# *De novo* RNA base editing in plant organelles with engineered synthetic P-type PPR editing factors

**DOI:** 10.1101/2024.09.13.612905

**Authors:** Sébastien Mathieu, Elena Lesch, Shahinez Garcia, Graindorge Stéfanie, Mareike Schallenberg-Rüdinger, Kamel Hammani

**Author notes:** These authors contributed equally (Sébastien Mathieu, Elena Lesch).

## Abstract

In plant mitochondria and chloroplasts, cytidine-to-uridine RNA editing plays a crucial role in regulating gene expression. While natural PLS-type PPR proteins are specialized in this process, synthetic PPR proteins offer significant potential for targeted RNA editing. In this study, we engineered chimeric editing factors by fusing synthetic P-type PPR guides with the DYW cytidine deaminase domain of a moss mitochondrial editing factor, PPR56. These designer PPR editors (dPPRe) elicited efficient and precise *de novo* RNA editing in *Escherichia coli*, and in *Nicotiana benthamiana* chloroplasts and mitochondria. Chloroplast transcriptome-wide analysis of the most efficient dPPRe revealed minimal off-target effects, with only three non-target C sites edited due to sequence similarity with the intended target. This study introduces a novel and precise method for RNA base editing in plant organelles, paving the way for new approaches in gene regulation applicable to plants and potentially other organisms.

---

RNA editing is a post-transcriptional process that introduces specific nucleotide changes in RNA sequences, which is essential for the proper functioning of many eukaryotic cells^1, 2^. As a result, it can alter the genetic information encoded in the DNA, allowing for the synthesis of proteins with amino acid sequences that differ from those dictated by the genomic template^3^. The ability to reprogram RNA editing specificity offers significant potential for biotechnological applications in basic research, medicine, and agriculture. It enables precise correction of genetic mutations, modulation of gene expression, regulation of protein function, and adaptation of cellular responses to various conditions.

In plants, RNA editing primarily involves the conversion of cytidines to uridines (C-to-U), and less frequently, uridines to cytidines (U-to-C) in organellar transcripts^4, 5, 6, 7, 8, 9^. In angiosperms like *Arabidopsis thaliana*, over 30 cytidines in chloroplast transcripts and more than 500 in mitochondrial transcripts are specifically edited^10, 11^. These modifications are crucial for the expression of functional proteins involved in essential metabolic and bioenergetic pathways^12, 13, 14, 15, 16, 17^. The specificity and efficiency of RNA editing in plants is mediated by nuclear-encoded pentatricopeptide repeat (PPR) proteins. PPR proteins are a class of RNA-binding proteins composed of tandem helix-turn-helix repeats of approximately 35 amino acids^18, 19, 20^. They are specific to eukaryotes and besides RNA editing, they regulate various post-transcriptional steps in organelles, from RNA stabilization to translation^21^. In plants, PPR proteins can be subdivided into two subfamilies based on the type and arrangement of their PPR arrays^22, 23^. P-type PPR proteins contain long stretches of canonical 35 amino acid P-PPR motifs and are widespread in eukaryotes, while PLS-type PPR proteins consist of arrays of P, L (long, 35-36 aa), and S (short, 31-32 aa) motifs and are specific to land plants performing organellar RNA editing^23, 24^. Genetic studies have implicated P-type PPR proteins in various post-transcriptional processes such as RNA stabilization, RNA cleavage, intron splicing, and mRNA translation^21^. In contrast, PLS-type PPR proteins are the key players in RNA editing^8^. They bind precisely to an RNA sequence located in the *cis*-element 4 nucleotides upstream of the target C to guide the deamination reaction (Fig. 1A). Each PPR motif (P, L, and S) aligns with a ribonucleotide in the *cis*-element in a parallel orientation and the RNA base recognition is mainly determined by a combinatorial code involving the identity of two amino acids in each P- and S-type PPR motifs: amino acid in position 5 distinguishes purines (A, G) from pyrimidines (C, U) and the last amino acid defines preferences for amino (A, C) or keto ribonucleobases (G, U)^25, 26, 27, 28^. The final P-L-S motif triplet (called P2-L2-S2) in PLS-PPR editing factors is often followed by E1 (for extension 1) and E2 motifs. The C-to-U conversion is catalyzed by a DYW deaminase domain of approximately 20 kDa, which is either naturally appended to the C-terminus of the PPR editing factor or recruited in *trans*^29, 30, 31, 32, 33, 34, 35^. Additionally, unusual SS-type PPR motifs were recently identified in several PLS-type PPR proteins of the lycophyte plant *Selaginella* and consists of tandem repeats of the S motif^22, 23^ (Fig. 1A).

**Fig. 1.**
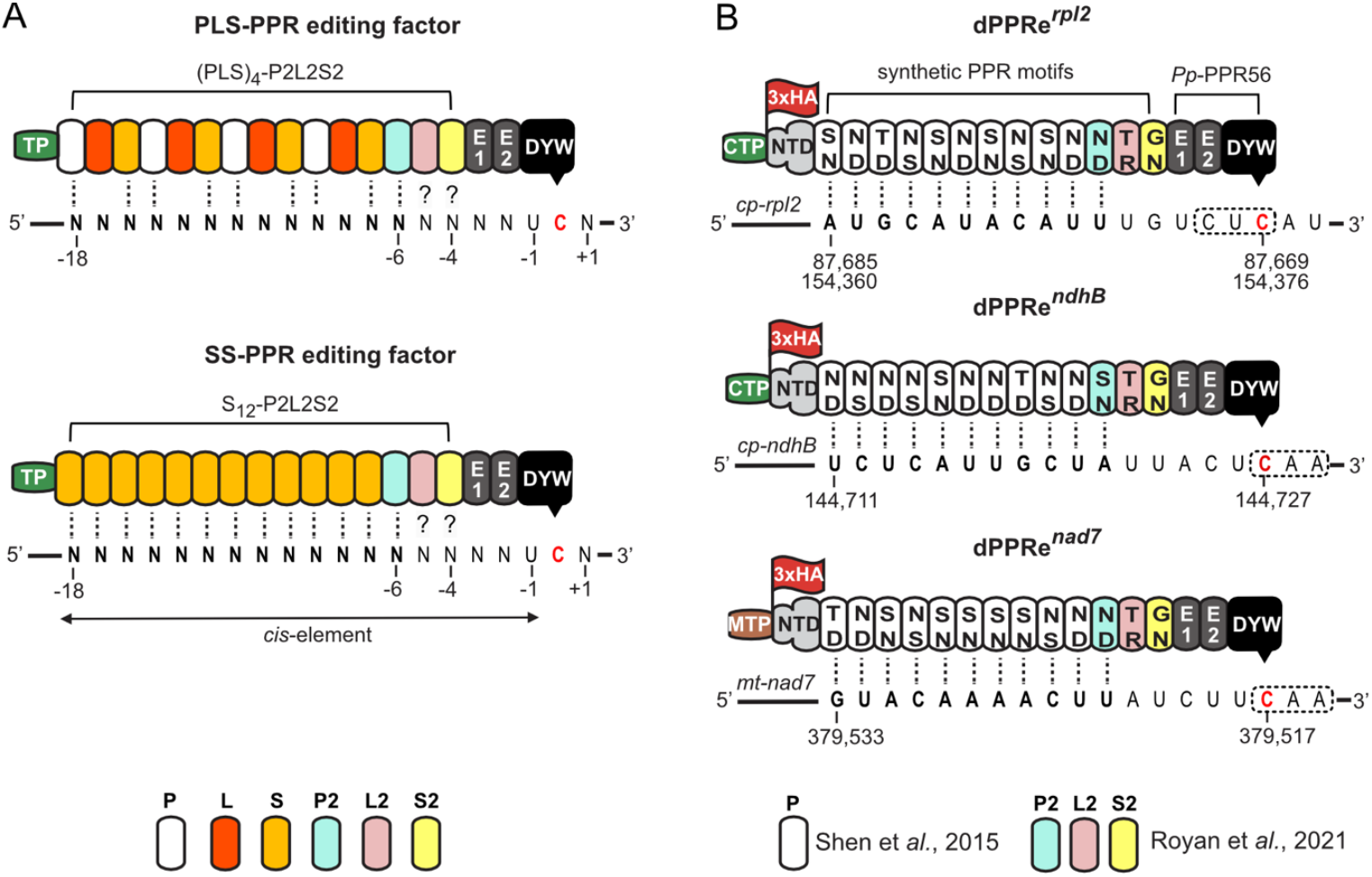
Design of precision RNA base PPR editors. **(A)** Motif composition and arrangement of natural PLS- and SS-type PPR editing factors in plant organelles. PPR motifs predicted to bind RNA bases are indicated by dotted black lines. Question marks indicate uncertain contribution to binding. The target cytidine is shown in red, and the designated PPR binding site (‘*cis*-element’) is highlighted in bold. The target cytidine is often preceded by a uridine in plant organellar genomes. TP: organellar transit peptide. **(B)** Engineering of chloroplastic and mitochondrial designer PPR editors (dPPRe) using arrays of synthetic P-type PPR motifs fused to the DYW cytidine deaminase domain from the mitochondrial editing factor PPR56. The specificity-determining amino acids at positions 5 and last of each PPR motif are indicated. The exact genomic position of the target cytidine in the *N. benthamiana rpl2* and *ndhB* chloroplast genes are shown beneath the RNA sequences, whereas the position within mitochondrial *nad7* gene was inferred from the mitochondrial genome sequence of *N. tabacum*. The codon containing the target cytidine is circled by a dotted box. CTP: chloroplast transit peptide, MTP: mitochondrial transit peptide. HA: hemagglutinin tag. NTD: N-terminal domain of PPR10 (amino acids 37 to 208).

The modular architecture of PPR proteins and the elucidation of the PPR base recognition code have enabled the rational design of synthetic PPR proteins using consensus P, L and S motifs with specific nucleotide binding capabilities^36, 37, 38, 39, 40^. *In vivo* applications of such synthetic P-PPR or PLS-PPR scaffolds for gene expression control in plants have been demonstrated in several contexts and include chloroplast RNA stabilization^41^, translational activation^42^, and RNA editing^39^. These studies have shown that synthetic PPR proteins can effectively mimic the functionality of their natural Arabidopsis counterparts in the respective mutant lines. Although these experiments provide proof-of-concept for synthetic PPR scaffolds being functional in chloroplasts, the potential to engineer these proteins for novel functions in gene expression by targeting new RNA sites in organelles has not yet been fully realized and requires further exploration. Additionally, the application of synthetic PPR scaffolds in mitochondria, which are associated with cytoplasmic male sterility in plants^43^ and inherited myopathies in humans^44^, remains largely unexplored.

In this study, we aimed to create *de novo* RNA editing in plant organelles by engineering chimeric editing factors. To this end, synthetic P-type PPR guides were fused to the DYW cytidine deaminase domain of a moss mitochondrial editing factor, PPR56^45^. We demonstrated that these designer PPR editors (dPPRe) function independently of cofactors and successfully edit their RNA targets in *E. coli*, as well as in the chloroplasts and mitochondria of *Nicotiana benthamiana* plants. Transcriptome-wide analysis of off-target effects of the most efficient dPPRe, in *N. benthamiana* chloroplasts with an editing efficiency up to 70% at its designated target C site in *rpl2* gene, revealed minimal off-target activity, affecting only three C sites with conserved *cis*-elements similar to the target site. This study demonstrates precise *de novo* RNA editing in plant organelles using engineered synthetic P-PPR repeats for the first time and provides a novel strategy for regulating gene expression by targeting RNA effectors, applicable to both chloroplasts and mitochondria in plants, and potentially other biological systems.

## Results

### Engineering synthetic *dPPRe* for precise RNA base editing

To date, two synthetic editing factors with PLS-type or SS-type PPR protein arrangements have been engineered to edit the chloroplast *rpoA*-78691 site *in vivo*, a site naturally targeted by the endogenous CLB19 PPR editing factor in Arabidopsis^36, 39^. The design of these factors involved concatenating synthetic domains derived from consensus P, L, S, P2, L2, S2, E1, E2, and DYW motifs to create protein scaffolds that mimic natural plant editing factors (Fig. 1A). Here, we adopted a different approach to design PPR editors (dPPRe) aimed at achieving precise *de novo* RNA editing in plant organelles. We hypothesized that any protein tract of sufficient length, with RNA-binding specificity toward the RNA sequence located six nucleotides upstream of a target cytidine (C), could effectively guide the DYW domain and catalyze cytidine deamination *in vivo*.

Unlike PLS-type PPR arrays, which contain L motifs whose contribution to RNA binding is unpredictable, we reasoned that P-type PPR arrays would be more suitable for reprogramming RNA editing specificity *in vivo*. This is due to their clear and predictable RNA-binding specificity, as all P motifs are involved in sequence recognition^25, 26^. To test this, we constructed guides composed of a tandem array of 10 synthetic P-type PPR motifs, termed designer PPR (dPPR)^37^ (Fig. 1B), which we and others have previously demonstrated to possess programmable RNA-binding specificity *in planta*^41, 42, 46^. A synthetic consensus P2-S2-L2 triplet was added after the last P motif to replicate the structural arrangement found in natural PPR editing factors. While the P2 motif is expected to confer RNA-binding specificity similar to P motifs, the roles of the L2 and S2 motifs in editing factors remain less clear^28, 47^. A guide composed of 11 P-type PPR repeats should be sufficient to specifically target an 11-nucleotide RNA sequence within plant chloroplast and mitochondrial genomes of few hundred kilobase pairs.

For the cytidine deamination reaction, we appended the C-terminal E1-E2-DYW domains from the *Physcomitrium patens* mitochondrial editing factor PPR56 to the synthetic [P]_10_-P2-L2-S2 scaffold. This particular E-DYW domain was chosen due to PPR56’s demonstrated ability to catalyze cytidine deamination across various biological environments, including plant mitochondria^45^ and cytoplasm^47^, *E. coli*^28, 33^, and human cell cytoplasm^48^. To stabilize the chimeric protein, an N-terminal domain (NTD) from the maize chloroplast-localized PPR protein PPR10^49^ was added upstream of the first P motif, resulting in the engineered dPPRe. Finally, we applied the PPR code to the [P]_10_-P2 motifs to target the RNA sequence located four nucleotides upstream of the selected cytidines, respectively.

For *in planta* editing assays, an N-terminal chloroplast transit peptide from Arabidopsis RecA (first 68 amino acids) or a mitochondrial presequence from yeast Cox4 (first 29 amino acids), both of which have been shown to efficiently direct fusion proteins to the respective organelles in plant cells^41, 50^, were added to the dPPRe constructs. A 3xHA tag was also included for immunodetection.

Initially, we engineered two chloroplast dPPRe constructs, dPPRe^*rpl2*^ and dPPRe^*ndhB*^, to target specific cytidines in the *rpl2* and *ndhB* mRNAs of *Nicotiana benthamiana*, whose chloroplast genome has been recently sequenced^51^ (Fig. 1B). These chloroplast genes encode a subunit of the large subunit of the chloroplast ribosome and the NADH dehydrogenase-like complex, respectively. Natively, these specific cytidines are not edited in the chloroplast transcriptome of *N. benthamiana*^51^. Editing the cytidine in the duplicated *rpl2* gene at positions 87,669 or 154,376 would affect the last base of the codon (CUC>CUU) without altering the amino acid sequence of the RPL2 protein (Leu>Leu) (Fig. 1B). In contrast, editing the cytidine at position 14,727 in *ndhB* mRNA is expected to change a CAA codon to UAA, introducing a premature stop codon and truncating the NdhB protein.

Before expressing the constructs *in planta*, we verified their functionality using a well-established heterologous *E. coli* RNA editing assay that allows co-expression of recombinant dPPRe (rdPPRe) without their transit peptide as N-terminal fusion proteins with a six-histidine and maltose-binding protein (MBP) tags, along with their RNA targets^33^ (Fig. 2). Upon isopropryl β-d-1thiogalactopyranoside (IPTG) induction of rdPPRe^*rpl2*^ and rdPPRe^*ndhB*^ expression in *E. coli*, a specific ∼130 kDa protein band corresponding to the expected molecular weight of the recombinant proteins accumulated in the bacterial total protein extracts (Fig. 2A). Sanger sequencing of cDNA from these extracts confirmed that rdPPRe successfully edited their target cytidines in *E. coli* (Fig. 2B and Table S1). Specifically, rdPPRe^*rpl2*^ edited its target cytidine with an efficiency ranging from 49% to 62% (average of 57% ± 6.8), while rdPPRe^*ndhB*^ exhibited a lower editing efficiency, averaging 33% ± 3.7. These results demonstrate that our chimeric dPPRe are functional for RNA editing *in vivo* and that synthetic P-type PPR arrays can effectively guide RNA editing by the DYW domain, similar to PLS- and S-type scaffolds.

**Fig. 2.**
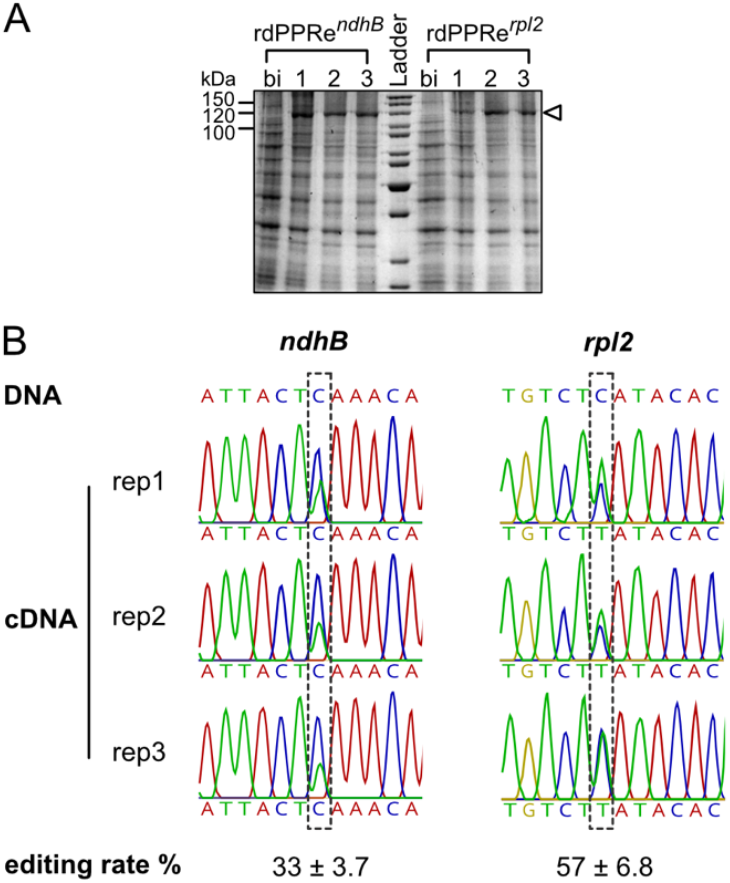
RNA editing activity of rdPPRe in *E. coli*. **(A)** Analysis of rdPPRe^*ndhB*^ and rdPPRe^*rpl2*^ protein expression in *E. coli* by SDS-PAGE and Coomassie blue staining of total protein extracts from three independent bacterial cultures harvested before induction (bi) or 20 hours after expression of constructs was induced. The arrowhead indicates protein bands whose apparent molecular weights correspond to the expected sizes for rdPPRe (131.5 kDa). **(B)** Sequencing electrophoregrams of cDNA from three independent *E. coli* cultures expressing rdPPRe^*ndhB*^ or rdPPRe^*rpl2*^ for 20 hours. Expression of rdPPRe in bacteria resulted in C-to-U editing at the co-expressed *rpl2* and *ndhB* target sites, as revealed by the appearance of a T peak in the cDNA sequence at these positions. The average editing rates are shown below (n=3).

### Expression of dPPRe in plants induces *de novo* RNA editing in chloroplasts

Next, we evaluated the RNA editing activities of dPPRe^*rpl2*^ and dPPRe^*ndhB*^ in chloroplasts through agroinfiltration-mediated transient gene expression in *Nicotiana benthamiana*. To enhance transient expression levels, dPPRe constructs were co-expressed with the tombusvirus suppressor of silencing, P19-HA^52^. Initially, we optimized assay conditions by testing *in vivo* protein accumulation of the most efficient dPPRe from *E. coli*, dPPRe^*rpl2*^, when driven by a single or double 35S promoter (2×35S), a strong and constitutive promoter, at 2 and 3 days post-infiltration (dpi). Immunodetection with anti-HA antibodies revealed that the 2×35S promoter facilitated higher accumulation of dPPRe^*rpl2*^ protein (expected molecular weight of ∼97 kDa) in *N. benthamiana* leaves compared to the single 35S promoter, with protein levels peaking at 3 dpi for both promoters (Fig. 3A).

**Fig. 3.**
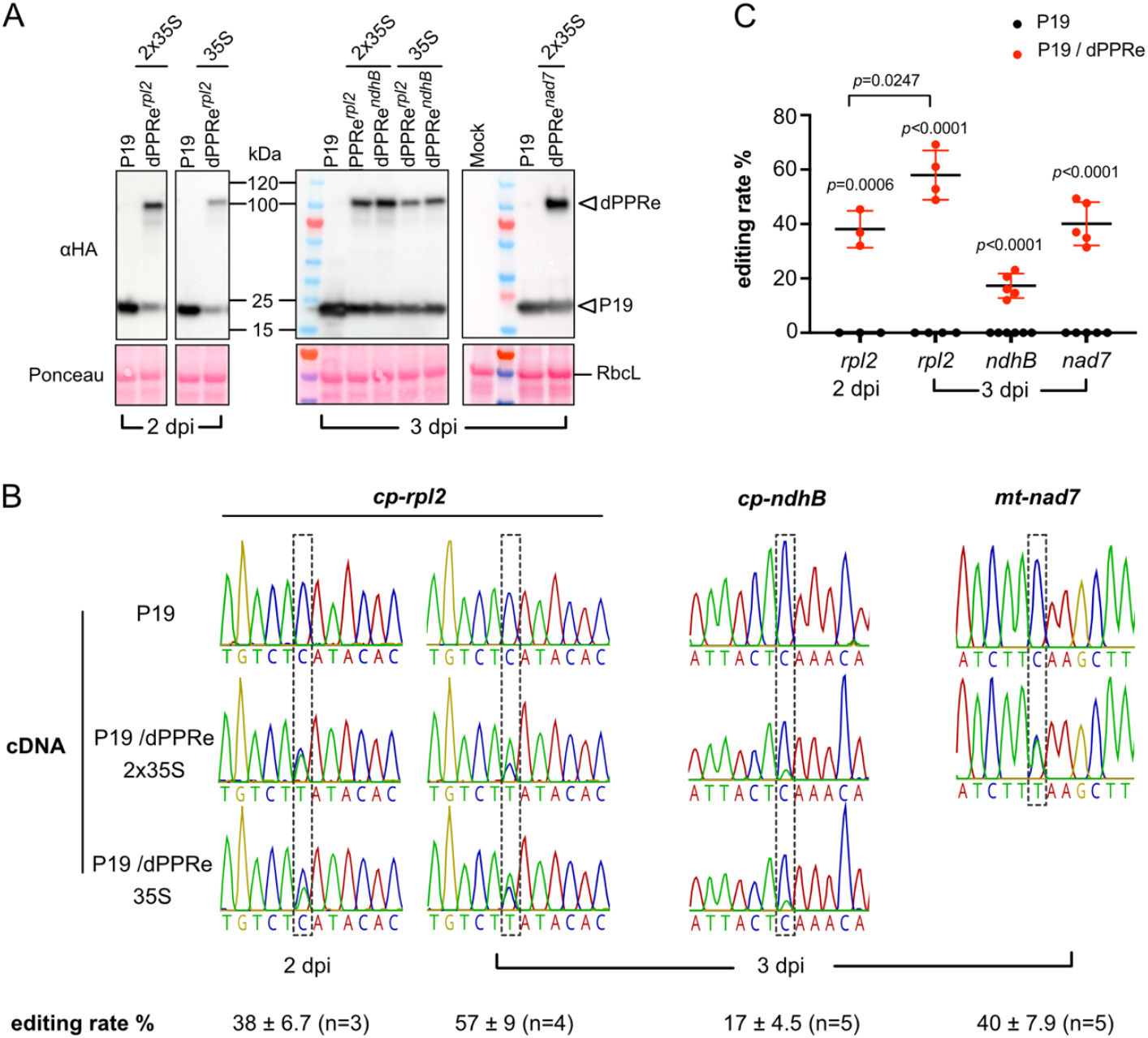
RNA editing activity of dPPRe *in planta*. **(A)** Immunodetection of chloroplast dPPRe^*rpl2*^, dPPRe^*ndhB*^ and mitochondrial dPPRe^*nad7*^ accumulation in *N. benthamiana* leaves at 2 and 3 days post-infiltration (dpi). Arrowheads indicate the protein bands corresponding to dPPRe (∼97 kDa) and the suppressor of silencing, P19 protein (∼19 kDa). **(B)** Sequencing electrophoregrams of cDNA from a replicate of each *in planta* experiment. Expression of dPPRe in *N. benthamiana* leaf led to *de novo* C-to-U RNA editing at the target sites in endogenous chloroplast *rpl2, ndhB* and mitochondrial *nad7* transcripts as revealed by the appearance of a T peak in the cDNA sequence at these positions that is absent in the control cDNA (P19 alone). The average editing rates with standard deviation (SD) and the number (n) of independently treated leaves are shown below (Table S2). **(C)** Editing rates at the respective targeted dPPRe sites for individual replicates from each experimental set up. For P19/dPPRe^*rpl2*^, the 2 dpi time point was assessed in three (n=3) independently treated *N. benthamiana* leaves, and the 3 dpi time point was assessed in four (n=4) leaves. The 3 dpi time point for P19/dPPRe^*ndhB*^ and P19/dPPRe^*nad7*^ was assessed in five (n=5) independently treated leaves. The P19 control for P19/dPPRe^*ndhB*^ at 3 dpi was assessed in six (n=6) leaves. Data are presented as means ± SD and were compared using *t*-tests with *p*-values shown in the graph (Table S2).

In parallel, we assessed RNA editing rates at the *rpl2*-87,669/154,376 sites in chloroplasts upon dPPRe^*rpl2*^ expression by cDNA sequencing. The results demonstrated that dPPRe^*rpl2*^ induced editing of the target cytidine in *rpl2* mRNA to an average of 38% at 2 dpi, increasing to 57% at 3 dpi while the expression of P19-HA alone did not cause editing (Fig. 3B-C and Table S2). A *t*-test confirmed the editing increase to be highly significant (*p*-value<0.05). Notably, editing efficiency at the target site could reach up to 69% at 3 dpi (Fig. 3C and Table S2). These findings suggest that dPPRe^*rpl2*^ functions as an effective RNA editing factor in plant chloroplasts, with higher activity observed at 3 dpi, indicating a doseable effect of dPPRe^*rpl2*^ on *rpl2* editing efficiency *in vivo*.

Subsequently, we expressed dPPRe^*ndhB*^ in *N. benthamiana* and examined RNA editing at *ndhB*-14,727 site at 3 dpi. Like dPPRe^*rpl2*^, dPPRe^*ndhB*^ showed higher protein accumulation with the 2×35S promoter (Fig. 3A). However, dPPRe^*ndhB*^ induced a lower editing rate *in planta*, achieving an average of 17% C-to-U conversion at its target site, reflecting the results obtained in *E. coli* (Fig. 3B-C, Table S2). This difference in editing efficiency between dPPRe^*rpl2*^ and dPPRe^*ndhB*^ in *E. coli* and *in planta* was not due to differences in protein accumulation (Fig. 1A-2A). Given that PPR protein binding to RNA can be inhibited by secondary structures^15, 53, 54^, we hypothesized that the stronger hairpin structure predicted at the dPPRe^*ndhB*^ binding site might impede RNA binding, leading to the reduced editing efficiency observed *in vivo* (Fig. S1).

Overall, these findings demonstrate that chimeric dPPRe proteins, composed of synthetic P-PPR guides and a mitochondrial E1-E2-DYW domain from moss, are functional in chloroplasts of angiosperms for *de novo* RNA editing. The results also suggest that the local folding of RNA target sequences plays an important role in determining dPPRe editing efficiency *in vivo*, consistent with previous reports for natural PPR editing factors^28, 55^.

### dPPRe^*rpl2*^ exhibits minimal off-target activity *in planta*

To evaluate the off-target RNA editing activity of dPPRe, we focused on the most efficient variant, dPPRe^*rpl2*^, and analyzed potential off-target sites in *N. benthamiana* chloroplasts. We performed RNA sequencing (RNA-seq) on samples from three independent agroinfiltration experiments. In these experiments, leaves were infiltrated either with the P19 protein alone (control) or with dPPRe^*rpl2*^/P19 driven by either a 35S or 2×35S promoter, resulting in nine cDNA libraries. To assess the relationship between *dPPRe*^*rpl2*^ expression levels and RNA editing rates, RNA was extracted at two time points: 2 dpi for one cDNA library and 3 dpi for two cDNA libraries in each of the three experiments.

Differential expression analysis using DESeq2 revealed no significant changes in chloroplast gene expression across the samples, with the exception of the *dPPRe*^*rpl2*^ nuclear transgene, which was significantly overexpressed in the experimental samples (Fig. 4A). This finding indicates that *dPPRe*^*rpl2*^ expression does not disrupt the overall regulation of chloroplast gene expression.

**Fig. 4.**
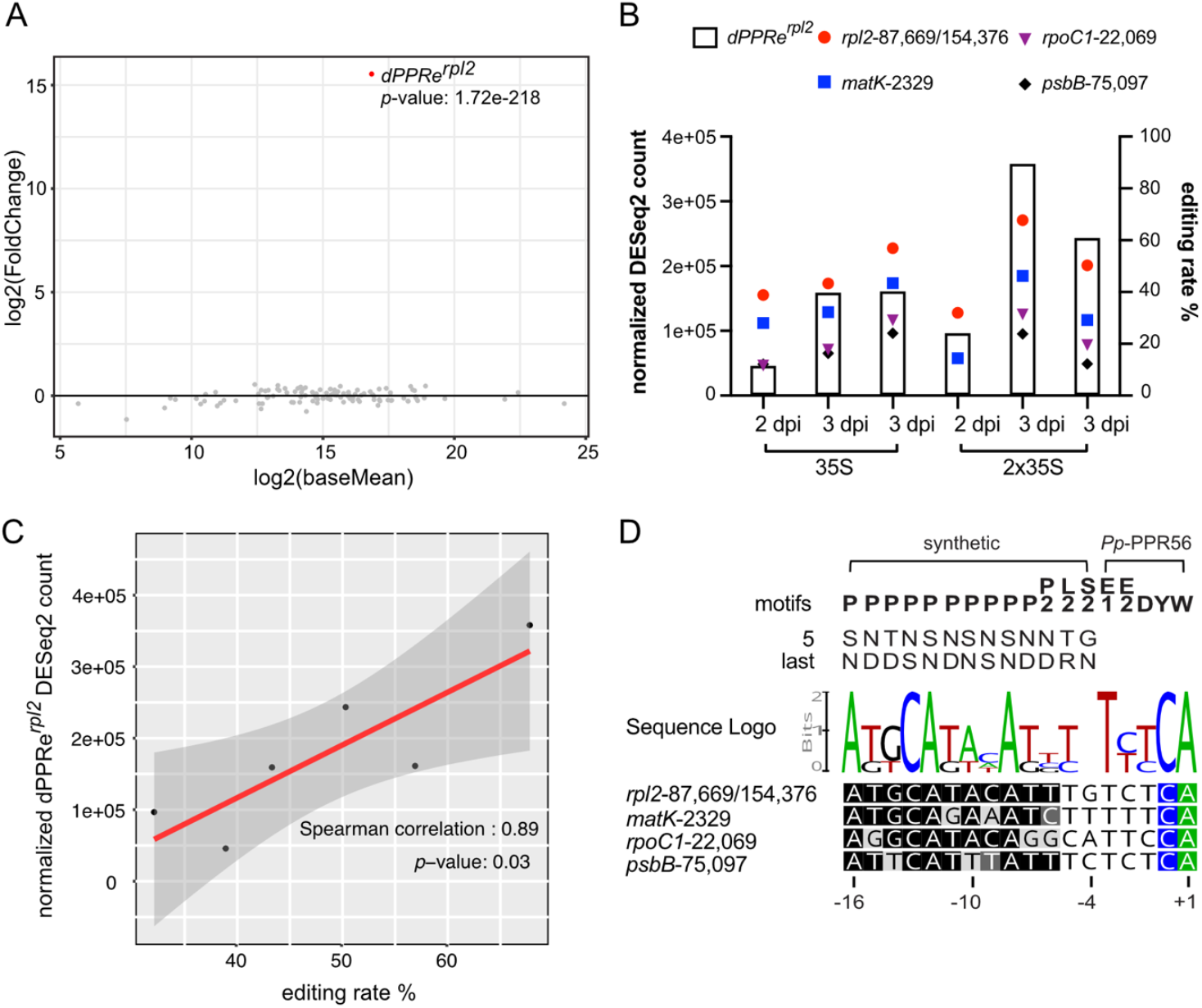
Off-target RNA editing activity of dPPRe^*rpl2*^ in *N. benthamiana* chloroplasts. **(A)** MA-plot displaying differentially expressed chloroplast genes and the *dPPRe*^*rpl2*^ transgene highlighted as a red dot. The plot displays log2 fold-changes in expression for dPPRe^*rpl2*^-expressing samples versus the normalized mean expression (log2 scale) for each gene. The dPPRe^*rpl2*^ transgene is the only gene with statistically significant differential expression (adjusted *p*-value < 0.05). **(B)** Bar plots depicting DESeq2-normalized dPPRe^*rpl2*^ expression in each experimental library, along with the corresponding C-to-U editing efficiency at identified *de novo* RNA editing sites. dpi: days post-inoculation. **(C)** Spearman correlation analysis between *dPPRe*^*rpl2*^ expression level and editing efficiency at *rpl2* target site. **(D)** Alignment between dPPRe^*rpl2*^ with its predicted binding sites upstream of the *de novo* chloroplast editing sites. A sequence logo was generated from the alignment of the *cis*-element sequences (−16/+1) at these sites. The PPR code determining amino acids five and last of each PPR motif are indicated, with nucleotide binding compatibility in the binding site sequences shaded from black to light gray, where light gray indicates no compatibility between the PPR code and the nucleotide in the binding site.

Next, we used the GATK4 toolkit to identify *de novo* C-to-U RNA editing sites triggered by dPPRe^*rpl2*^ expression (Table S2). As expected, the target cytidine in the *rpl2* gene was edited (Fig. 4B). However, we also detected three off-target editing sites in the *matK, rpoC1*, and *psbB* chloroplast genes. Editing at the *rpl2* target site and the *matK-*2329 site was consistently observed across all *dPPRe*^*rpl2*^-expressing libraries, while editing at the *rpoC1-*22,069 and *psbB-*75,097 sites was detected in only five of the six libraries (Fig. 4B).

We quantified the editing rate at these four sites in each experimental library in relation to the expression levels of *dPPRe*^*rpl2*^ (Fig. 4B). Consistent with previous protein immunodetection results, the 2×35S promoter drove higher expression of *dPPRe*^*rpl2*^ than the single 35S promoter, and *dPPRe*^*rpl2*^ expression was higher at 3 dpi compared to 2 dpi for both promoters. The quantification revealed that the *rpl2* target site had the highest editing rate (ER) across all libraries, reaching up to 67.8%, followed by *matK*-2329 (ER ≤ 46.3%), *rpoC1*-22,069 (ER ≤ 31.4%), and *psbB*-75,097 (ER ≤ 21.9%) (Fig. 4B). These results suggest that dPPRe^*rpl2*^ preferentially guides RNA editing towards its intended cytidine target over off-target sites *in vivo*. Moreover, comparison of editing rates across independent experiments indicated a positive correlation between *dPPRe*^*rpl2*^ expression levels and RNA editing efficiency (Fig. 4B), particularly at the *rpl2* target site, as confirmed by a Spearman correlation test (*p*-value < 0.05) (Fig. 4C and Fig. S2).

To further investigate the off-target activity of dPPRe^*rpl2*^, we aligned the *cis*-element sequences spanning from nucleotide -16 to +1 relative to each off-target site with that of the *rpl2* target site. This analysis revealed a conserved nucleotide sequence among the off-target and *rpl2 cis*-elements (Fig. 4D). Notably, the first 11 nucleotides, predicted to be bound by the synthetic [P]_10_-P2 guide, exhibited high conservation, with all off-target sites sharing 8 nucleotides with the *rpl2* target site within this region. Additionally, a T was observed at position -3, and an A immediately downstream of the edited cytidine in all off-targets.

Further analysis of the nucleotide preferences of the 11 PPRs in dPPRe^*rpl2*^ showed that some motifs tolerate mismatches along the 11-nucleotide target sequence (Fig. 4D). The number and position of these mismatches within the PPR-RNA duplex appeared to influence *in vivo* editing efficiency, likely by affecting protein-RNA interactions. For example, the most efficiently edited off-target, *matK*-2329, contained two mismatches positioned one nucleotide apart in the central region of the dPPR binding site. In contrast, the less efficiently edited sites, *rpoC1*-22,069 and *psbB*-75,097, exhibited two to three mismatches distributed differently: *rpoC1*-22,069 had three mismatches spread across the 5’ and 3’ ends of the RNA, while *psbB-75,097* had two mismatches located between the 5’ end and the middle of the RNA. These findings suggest that the dPPR guide is more tolerant of mismatches in the central region of its binding site than at the 5’ or 3’ ends.

### dPPR editors are active in plant mitochondria

Finally, we evaluated the potential of dPPRe for *de novo* RNA editing in plant mitochondria. For this purpose, we engineered dPPRe^*nad7*^ to edit a specific cytidine in the *nad7* mRNA within *N. benthamiana* mitochondria, which encodes a subunit of the respiratory complex I (Fig. 1B). Since the mitochondrial genome of *N. benthamiana* has not yet been sequenced, we selected this editing site based on the sequence of the mitochondrial *nad7* gene from *Nicotiana tabacum*, assuming that the sequence is conserved between the two species. Similar to the editing event at *ndhB*-14,727, RNA editing at the selected cytidine in the mitochondrial *nad7* gene is expected to create a premature stop codon in the mRNA (CAA>UAA) (Fig. 1B).

dPPRe^*nad7*^ was expressed under the control of a 2×35S promoter in *N. benthamiana* leaves and successfully accumulated by 3 dpi, as confirmed by anti-HA immunodetection. cDNA sequencing confirmed *nad7* sequence and revealed that the expression of dPPRe^*nad7*^ in plant cells induced RNA editing at the target cytidine within mitochondria, achieving an ER of up to 49% (Fig. 3B-C). This result demonstrates that synthetic dPPRe are functional ribocytidine editors in plant mitochondria.

## Discussion

The results of this study highlight the significant potential of synthetic designer PPR RNA editors, as a powerful tool for precise RNA base editing in plant organelles, with successful applications demonstrated in both chloroplasts and mitochondria. Our engineering of dPPRe constructs, which incorporate arrays of synthetic P-type PPR motifs fused with the DYW cytidine deaminase domain from the moss PPR56 editing factor, represents a significant advancement over previous designs that relied on PLS-type or SS-type PPR arrangements.

Previously, the first synthetic PLS-PPR editing factor (referred to as dsnPLS^CLB19^) was capable to substitute the function of the natural CLB19 PLS-PPR editing factor at the *rpoA*-78,691 site in the respective Arabidopsis knockout line^39^. However, efficient editing was only possible in the presence of plant protein cofactors from the RIP/MORF family (Multiple Organellar RNA editing Factors). Structural and biochemical studies suggest that MORF interaction with PLS-PPR editing factors, particularly with the L motifs, enhances RNA-binding efficiency by inducing conformational changes^56^. Consequently, the application of synthetic PLS-PPR scaffolds may be limited to angiosperm plants that possess MORF proteins^57, 58^. Moreover, despite higher transgene expression levels, dsnPLS^CLB19^ achieved only ∼45% editing efficiency *in vivo* compared to over 80% efficiency by the natural CLB19 editing factor, highlighting its limited effectiveness compared to its natural counterpart.

To overcome the MORF-dependency, a subsequent SS-type PPR scaffold (dsnS^CLB19^) was developed using a similar approach. dsnS^CLB19^ edited its RNA target in *E. coli* without plant MORF cofactors, achieving an editing rate below 50%, similar to dsnPLS^CLB1936^. However, the effectiveness of the dsnS scaffold *in planta* remains unconfirmed, and the broader applicability of dsnPLS- and dsnS-type editors for reprogramming RNA editing specificity in plant organelles is still uncertain. In contrast, our study demonstrates that synthetic P-type dPPR editors can induce *de novo*, site-specific RNA editing in both chloroplasts and mitochondria achieving high efficiency of up to 69%. This significantly expands the RNA editing toolkit for plant systems.

The DYW deaminase of PPR56 has been shown to function in various biological systems including human cells, plants and bacteria without perturbating the target cell^33, 45, 48^. In this study, we demonstrated that when fused to a programmable P-type PPR array, this deaminase can rewrite the genetic information within a target RNA by enzymatically converting a specific cytidine in both plant chloroplasts and mitochondria. This makes dPPRe a versatile tool for site-directed RNA base editing. However, like other site-directed RNA editing approaches, dPPRe face challenges related to editing efficiency and target scope. Although dPPRe can achieve a satisfactory RNA editing efficiency over 50% for biotechnological applications, certain plant PPR editing factors, such as the native PPR56, can nearly fully convert their cytidine target *in vivo*^33, 45^. The lower efficiency of the chimeric dPPRe may be due to the E1-E2-DYW domains of PPR56 working in concert with the PPR array for RNA recognition and editing^59, 60^, a cooperation that might not be fully supported by synthetic PPR scaffolds, leading to reduced *in vivo* activity.

In addition, similar to ADAR (Adenosine Deaminases Acting on RNA) and APOBEC (Apolipoprotein B Editing Catalytic polypeptide) base editors^61, 62, 63^, the DYW deaminases of natural PPR editing factors and their associated E1-E2 domains have nucleotide context preferences, preferring cytidine that are embedded in specific RNA sequences^59, 60, 64^. For instance, the -3/+1 nucleotide sequences surrounding the dPPRe^*rpl2*^ off-target cytidines in chloroplasts are rich in U/C and A (Fig. 4D), aligning with the described preferences of PPR56^47, 59, 64^. This nucleotide context preference might also explain the lower editing efficiency of dPPRe^*ndhB*^ *in vivo*: sequence analysis of dPPRe^*rpl2*^ off-targets revealed a strict preference of PPR56’s E1-E2-DYW domain for U at position -3, while the *ndhB* target C site has an A at this position (Fig. 1B). While this nucleotide preference is beneficial for the engineering of site-specific RNA editors by increasing editing precision, it limits the number of editable cytidines in a given transcriptome.

Furthermore, this study provides important insights into the precision and efficiency of RNA editing achieved by dPPRe *in planta*. While dPPRe^*rpl2*^ exhibited robust editing rates at its target site in the *rpl2* gene *in planta*, we also detected a few off-targets. These off-target sites, however, shared significant sequence similarities with the intended target, particularly in the regions predicted to interact with the synthetic PPR motifs. Our findings underscore the importance of carefully designing PPR motifs to minimize unintended interactions, as PPR-RNA interactions are not solely defined by the PPR RNA-binding code itself^25^. For instance, when we applied the PPR code to predict binding sites for the [P]_10_-P2 motifs of dPPRe^*rpl2*^ and possible cytidine targets within the chloroplast genome of *N. benthamiana*, the designated cytidine target in *rpl2* was identified as the top hit (Fig. S3). However, none of the other predicted targets were edited in plants expressing dPPRe^*rpl2*^, and the three off-targets observed *in planta* were not among the top 40 matches. This discrepancy lies in the difficulty to evaluate the energetic costs of mismatches and their positional effects within a PPR-RNA duplex^38^, competing RNA secondary structures^38^, and the influence of the C-terminal DYW domains on nucleotide specificity^65^}^47, 59, 64^. A deeper understanding of these effects will significantly enhance our ability to predict PPR-RNA interactions and improve the specificity of dPPRe design without compromising editing efficiency in the future.

Recently, the use of the *E. coli* editing system and a plant cytosolic editing assay, demonstrated that the DYW domains from plant editing factors besides PPR56 show some variations in their flexibility towards the nucleotide context around the cytidine target site^47, 64^. These DYW domains offer great promise for broadening the editing site-specific capability of dPPRe. Although synthetic E1, E2, and DYW domains engineered from consensus amino acid sequences are currently available^39^, their preferred nucleotide context remains unknown and complicates their use for *de novo* RNA editing *in vivo*.

In conclusion, the successful application of dPPRe in *E*.*coli*, chloroplasts and mitochondria emphasizes their versatility for targeted RNA editing and provides the first mitochondrial RNA base editor tool kit. Mitochondrial dysfunction is associated with a range of phenotypic abnormalities in plants, including cytoplasmic male sterility, which is of considerable interest in hybrid seed production^66^. The ability to precisely edit mitochondrial genes could therefore have profound implications for the development of new plant varieties with improved traits. Beyond plant systems, dPPRe hold great promise for mitochondrial research and associated human diseases^44^. Given that PPR56 is active for RNA editing in human cells^48^, dPPRe technology could pave the way for innovative therapeutic strategies targeting mitochondrial genes. As we continue to refine and expand the capabilities of dPPRe, this technology holds the potential to become a versatile and indispensable tool in organellar genetic research.

## Methods

Primers used in this study are listed in Table S3 and DNA and protein sequences are provided in Figure S4.

### *dPPRe* transgene cloning

The DNA sequences encoding the Arabidopsis chloroplast RecA or yeast mitochondrial Cox4 transit peptide (68 and 29 amino acids, respectively) fused to a 3xHA tag and dPPRe were synthesized and cloned into pDONR™/Zeo by Thermo Fisher Scientific. For plant expression, the genes were subsequently cloned into binary vectors pMDC32 (2×35S promoter)^67^ or pGWB2 (35S promoter)^68^ using LR clonase II (Thermo Fisher Scientific, Waltham, MA, USA) following the manufacturer’s protocol.

For expression in *Escherichia coli*, the coding sequences of mature dPPRe without the organellar transit peptide and 3xHA tag were amplified from the pDONR™/Zeo templates. Target sequences comprising 100 bp up- and downstream of the cytidine to be edited, respectively, were amplified from *Nicotiana benthamiana* genomic DNA. dPPRe and RNA target encoding sequences were combined via Overlap Extension PCR and inserted into petG41K_MCS using FastDigest restriction endonucleases KpnI and SgsI (Thermo Fisher Scientific).

### Bacterial RNA editing assays

*E. coli* RNA editing assays were performed as outlined in Oldenkott et al., 2019. In short, constructs were transformed into *E. coli* Rosetta 2(DE3) competent cells (Novagen), which were grown in liquid culture until reaching an optical density (OD_600_) of 0.4-0.6 before inducing recombinant dPPRe (rdPPRe) expression by the addition of 0.4 mM IPTG. Cells were harvested after 20 hours of incubation at 16°C under continuous shaking. RNA editing was determined by RT-PCR and Sanger sequencing and rdPPRe expression was verified by SDS-PAGE and Coomassie blue staining. The mean RNA editing rates were calculated from three biological replicates (independent primary *E. coli* clones that underwent individual RNA editing assays).

### Transient expression in *Nicotiana benthamiana*

For transient protein expression, 6-week *Nicotiana benthamiana* plants were infiltrated with *Agrobacterium tumefaciens* GV3101 carrying the *dPPRe* transgenes. Agrobacterium cultures were grown overnight, resuspended in infiltration buffer (10 mM MgCl_2_, 10 mM MES, pH 5.6, and 100 μM acetosyringone) to an OD_600_ of 0.7, and infiltrated into the abaxial side of fully expanded *N. benthamiana* leaves using a needleless syringe. A Tomato bushy stunt virus-encoded 35S:*P19-HA* silencing suppressor gene construct^52^ (P19) was co-expressed with *dPPRe* transgenes to increase protein expression. Mixtures of P19/dPPRe agrobacteria were prepared in a 1:1 (v/v) ratio as required. P19 alone was used as the control. Infiltrated plants were maintained under controlled growth conditions and leaf discs were harvested at 2- and 3-day post-infiltration (dpi) for protein analysis by Western Blot using monoclonal anti-HA antibody (H9658 clone) from Sigma-Aldrich. Entire leaves were also collected at the same time points for RNA extraction using the TRIzol reagent, following the manufacturer’s protocol, to assess RNA editing and gene expression levels by sanger sequencing of RT-PCR products or RNA-seq.

### Plant RNA sequencing

Total RNA was extracted from *N. benthamiana* leaves infiltrated with dPPRe^*rpl2*^ and P19 constructs or P19 alone at 2-dpi or 3-dpi by Trizol extraction followed by phenol/chloroform extraction and ethanol precipitation. RNA was treated by DNase Turbo (Thermo Fisher Scientific) to remove contaminant DNA. Library preparation (rRNA-depleted) and 2×150-bp pair-ended Illumina sequencing were done by Novogene (https://www.novogene.com/), generating ∼105-138 mio reads per library. Libraries were prepared and sequenced for three independent biological replicates of P19, dPPRe^*rpl2*^ (2×35S)/P19 and dPPRe^*rpl2*^ (35S)/P19 transient expression.

The quality of the reads was assessed with FastQC 0.12.1 and adapters were trimmed with Trim-galore 0.6.10. Sequences with a quality lower than 30, with more than 5 ‘N’s or shorter than 20 bp after trimming were removed. The remaining reads were mapped against the *N. benthamiana* chloroplast genome SRR7540368^51^ (https://github.com/WangSoybean/Nicotiana_chloroplast_genomes) combined with the *dPPR*^*rpl2*^ nucleotide sequence with HISAT2 2.2.1 using the -k 50 option and default parameters. Mapped reads were counted with featureCounts 2.0.1 with options -O -M --minOverlap 5, to consider multi-mapped reads. Differential expression analysis was done using the DESeq2 1.42.0 package.

Chloroplast RNA editing sites were identified from the HISAT2 aligned reads with GATK 4.5.0.0 according to the GATK best practice recommendations^69, 70^ (https://gencore.bio.nyu.edu/variant-calling-pipeline-gatk4/). Initially, MarkDuplicatesSpark, HaplotypeCaller, and SelectVariants were run with default parameters, and VariantFiltration with options “QD_filter” -filter “QD < 2.0”, -filter-name “FS_filter” -filter “FS > 60.0”, -filter-name “MQ_filter” -filter “MQ < 40.0”, -filter-name “SOR_filter” -filter “SOR > 4.0”, -filter-name “MQRankSum_filter” -filter “MQRankSum < -12.5”, -filter-name “ReadPosRankSum_filter”, -filter “ReadPosRankSum < -8.0”. This initial run served as input for BaseRecalibrator, ApplyBQSR, and BaseRecalibrator steps. These steps were then repeated with the same options to call single nucleotide polymorphisms (SNPs) between the chloroplast transcriptome and genome. RNA editing sites were identified by examining C-to-T or G-to-A conversions. *De novo* RNA editing sites induced by dPPRe^*rpl2*^ expression were identified by detecting base conversions specifically present in experimental libraries (dPPRe^*rpl*2^/P19) but absent in control libraries (P19). The editing sites introduced by the expression of dPPRe^*rpl2*^ were considered significant if they were detected in at least two independent libraries. Editing rates were calculated as the ratio between the number of SNP reads and read depth at the specific base position. The spearman correlation test (cor.test in R 4.2.0) was used to assess the correlation between the editing rate and expression level of dPPRe^*rpl2*^.

The MA-plot and correlation test graphs were generated using the ggplot2 3.5.1 R package. Secondary structures were predicted with the mfold server (RNA folding form version 2.3) at http://www.unafold.org/ using default parameters and a folding temperature of 24°C.

### Statistics and reproducibility

All experiments were repeated at least 3 times. The number of samples for each experience is provided in the figures. The statistical significance of the RNA editing rate differences between experimental and control groups or between experiments in Figure 4B was assessed by *t*-test using GraphPad Prism 8.4.0 (*t*-values and degrees of freedom can be found in Table S2).

## Data availability

The read data were deposited to the NCBI Sequence Read Archive database with the Bioproject identifier GSEXXXXX.

## Acknowledgements

This work was supported by Agence National de la Recherche [ANR-18-CE20-0013 to K.H.] and Centre National de la Recherche Scientifique. Work of M.S-R. and E. L was funded by the Deutsche Forschungsgemeinschaft (grant: SCHA 1952/2-2). The authors thank Volker Knoop for useful discussions.

## Authors contributions

S.M., E.L., M.S.-R. and K.H. designed and conceived the experiments. E.L. performed the *E. coli* experiments. S.M., Sh.G., and K.H. performed the *in planta* work. S.G. performed bioinformatic analysis. S.M., E.L., S.G. and K.H. analyzed the results. K.H. supervised the work, made the figures and drafted the manuscript. S.M., E.L., M.S.-R. edited the manuscript.

## Supplemental information

**Fig. S1**. Mfold prediction of the most stable structure of the -100/+100 ribonucleotide sequences surrounding the cytidine targeted by dPPRe *in vivo*. The predicted dPPRe binding sites are underlined in blue, while the target cytidine is circled and marked with an arrowhead. Free energy values (Δ*G*) are provided for both the full RNA structures and the local RNA structures containing the dPPRe binding sites (indicated by a dotted line box).

**Fig. S2**. Spearman correlations between dPPRerpl2 expression levels, as determined by DESeq2, and off-target editing rates.

**Fig. S3**. Weight matrix used for prediction of dPPRe^*rpl2*^ cytidine targets within the *N. benthamiana* chloroplast genome by the TargetScan tool of PREPACT^71^ and the scanning output with the 40 best predicted targets. (*Top*) Each column corresponds to one nucleotide position and is labelled either (i) with type and fifth and last amino acid of the corresponding PPR or (ii) with the position relative to the editing site (EdS). Nucleotide distributions were set according to the published PPR-RNA binding code. Positional weighs are shown below. (*Bottom*) Best 40 predicted targets with the position in the genome and the matrix score (maximum 1000). Matches are colored in green and mismatches in red and given are the percentages of the match.

**Fig. S4**. Sequences of dPPRe^*rpl2*^, dPPRe^*ndhB*^ and dPPRe^*nad7*^. Arabidopsis RecA and yeast Cox4 transit peptides are in bold black, the 3xHA tag is shown in red and the synthetic P-PPR array is delimited by a box. The C-terminal synthetic P2, S2, L2 and PPR56’s E1-E2-DYW are highlighted in turquoise, pink, yellow and grey, respectively. The starting amino acid of the dPPR protein expressed as a recombinant protein in *E. coli* is shaded in red.

**Table S1**. Editing rates of rdPPRe in *E. coli*

**Table S2**. Editing rates of dPPRe *in planta*

**Table S3**. Differential gene expression analysis (DEseq2) data used for MA-plotting

**Table S4**. GATK analysis

**Table S5**. Oligonucleotide sequences used in this study

