## Supplemental file for "*De novo* RNA base editing in plant organelles with engineered synthetic P-type PPR editing factors"

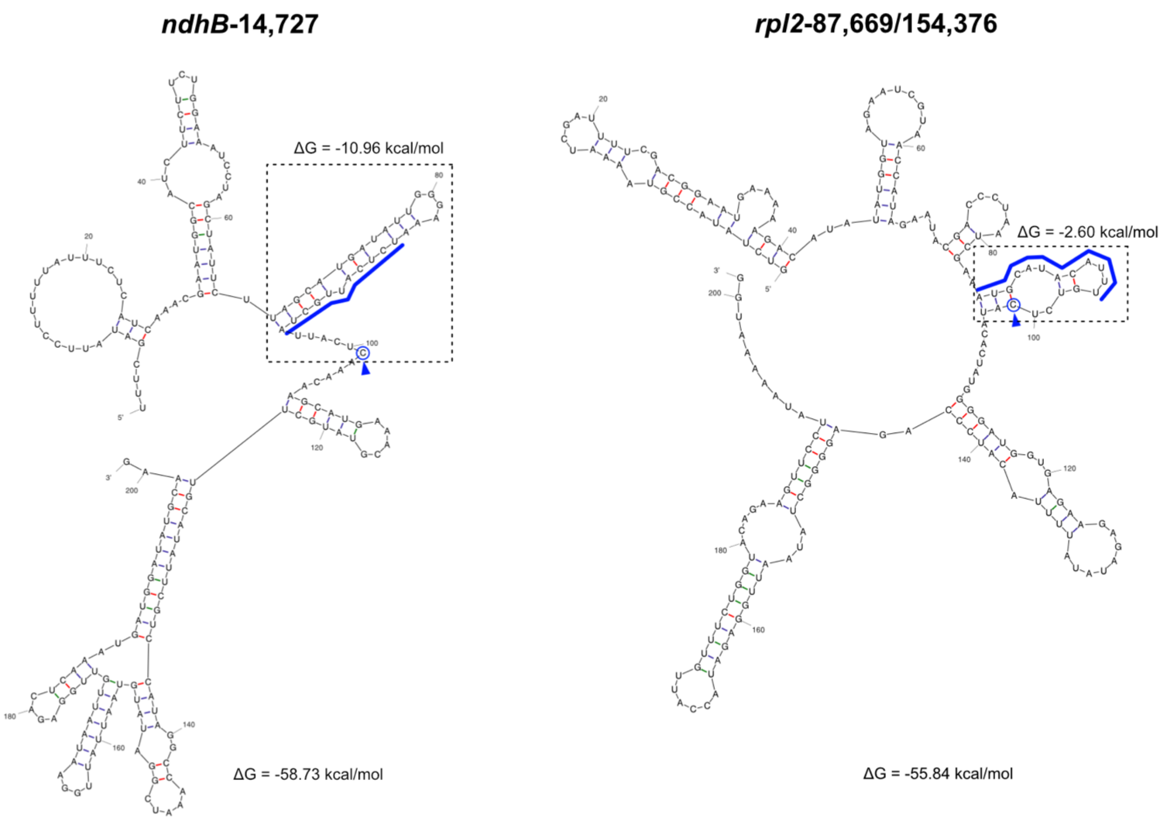


**Fig. S1.** Mfold prediction of the most stable structure of the -100/+100 ribonucleotide sequences surrounding the cytidine targeted by dPPRe *in vivo*. The predicted dPPRe binding sites are underlined in blue, while the target cytidine is circled and marked with an arrowhead. Free energy values (Δ*G*) are provided for both the full RNA structures and the local RNA structures containing the dPPRe binding sites (indicated by a dotted line box).

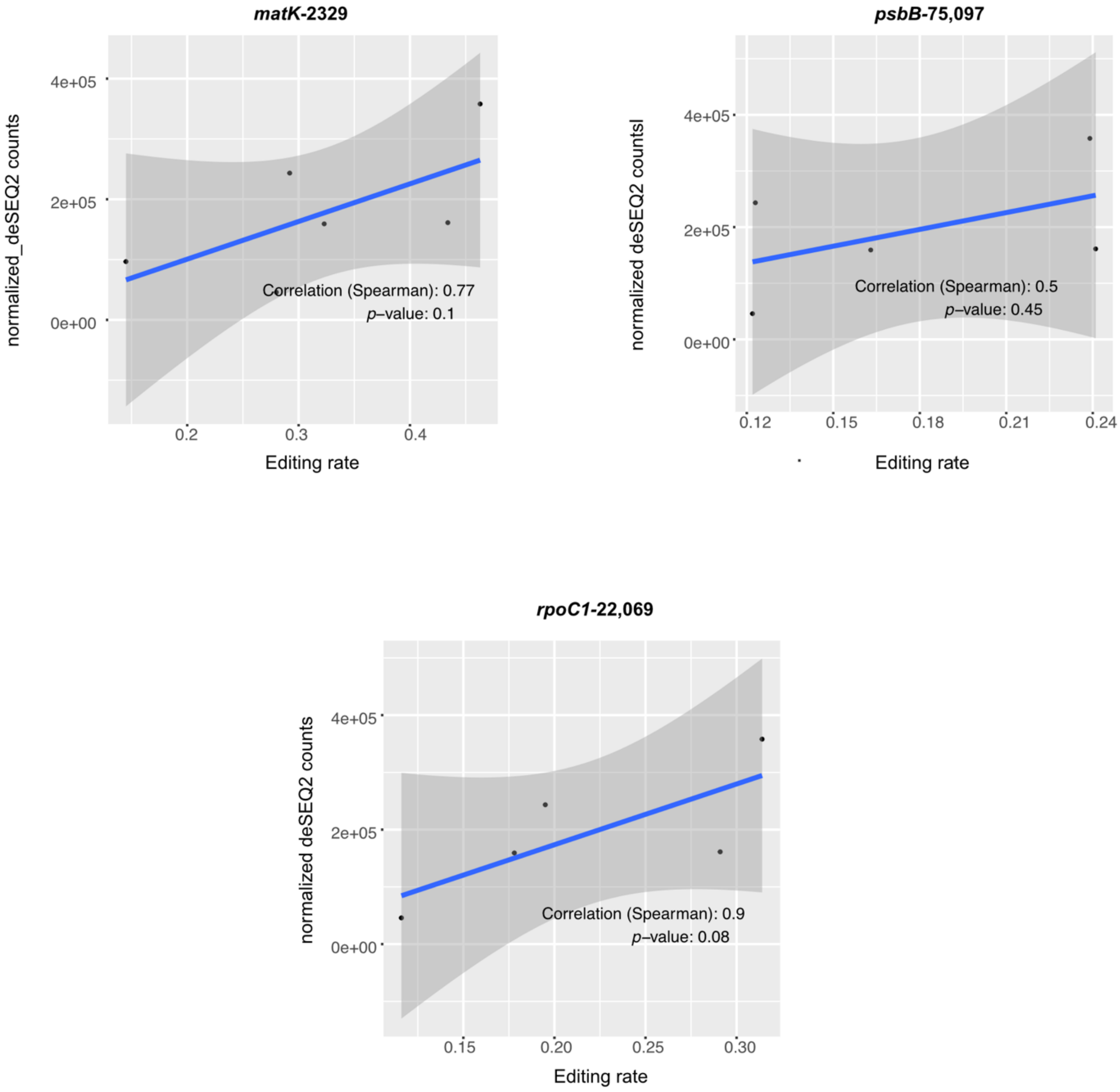


**Fig. S2.** Spearman correlations between dPPRe$^{rpl2}$ expression levels, as determined by DESeq2, and off-target editing rates.

### Sequence weight matrix

**Matrix settings**

18 base positions ?

**File handling**

Upload matrix data ? Download matrix data ?

Matrix to adjust positional base composition and weighting (see help for recalculation and locking): ?

5' end

| | P:SN | P:ND | P:TD | P:NS | P:SN | P:ND | P:SN | P:NS | P:SN | P:ND |
|---|---|---|---|---|---|---|---|---|---|---|
| % A | 80 | 0 | 20 | 0 | 80 | 0 | 80 | 0 | 80 | 0 |
| % C | 0 | 40 | 0 | 60 | 0 | 40 | 0 | 60 | 0 | 40 |
| % G | 20 | 0 | 80 | 0 | 20 | 0 | 20 | 0 | 20 | 0 |
| % T | 0 | 60 | 0 | 40 | 0 | 60 | 0 | 40 | 0 | 60 |
| x % | 100 | 100 | 100 | 100 | 100 | 100 | 100 | 100 | 100 | 100 |

| | P2:ND | L2 | S2 | -3 | -2 | -1 | EdS | +1 |
|---|---|---|---|---|---|---|---|---|
| % A | 0 | 25 | 25 | 25 | 25 | 15 | 0 | 25 |
| % C | 40 | 25 | 25 | 25 | 25 | 40 | 100 | 25 |
| % G | 0 | 25 | 25 | 25 | 25 | 5 | 0 | 25 |
| % T | 60 | 25 | 25 | 25 | 25 | 40 | 0 | 25 |
| x % | 100 | 0 | 0 | 0 | 0 | 100 | 200 | 0 |

3' end

**Top 40 hits**

| No. | Accession | Taxon | Organelle | Search result location | Scores/sequence ? | Total (max. 1000) |
|---|---|---|---|---|---|---|
| 1 | SRR7540368 | | | 154360..154377 | 80 60 80 60 80 60 80 60 80 60 60 0 0 0 0 40 200 0<br>A T G C A T A C A T T T G T C T C A | 1000 |
| 2 | SRR7540368 | | | complement(87668..87685) | 80 60 80 60 80 60 80 60 80 60 60 0 0 0 0 40 200 0<br>A T G C A T A C A T T T G T C T C A | 1000 |
| 3 | SRR7540368 | | | 1882..1899 | 80 60 80 40 80 60 80 60 80 60 40 0 0 0 0 40 200 0<br>A T G T A T A C A T C A G T T C C T | 960 |
| 4 | SRR7540368 | | | 73416..73433 | 80 60 80 40 80 60 80 0 80 60 60 0 0 0 0 40 200 0<br>A T G T A T A G A T T C C A G T C C | 920 |
| 5 | SRR7540368 | | | 20500..20517 | 80 60 80 40 80 60 80 40 80 60 40 0 0 0 0 5 200 0<br>A T G T A T A T A T C A T C T G C T | 905 |
| 6 | SRR7540368 | | | 81678..81695 | 80 60 80 0 80 60 80 40 80 60 40 0 0 0 0 40 200 0<br>A T G G A T A T A T C C A T T T C A | 900 |
| 7 | SRR7540368 | | | 99209..99226 | 80 60 80 40 80 60 80 0 80 60 40 0 0 0 0 40 200 0<br>A T G T A T A G A T C C T G T T C A | 900 |
| 8 | SRR7540368 | | | 122261..122278 | 80 40 20 40 80 60 80 60 80 60 60 0 0 0 0 40 200 0<br>A C A T A T A C A T T T C A A C C T | 900 |
| 9 | SRR7540368 | | | complement(142819..142836) | 80 60 80 40 80 60 80 0 80 60 40 0 0 0 0 40 200 0<br>A T G T A T A G A T C C T G T T C A | 900 |
| 10 | SRR7540368 | | | 2273..2290 | 80 0 80 40 80 60 80 40 80 40 60 0 0 0 0 40 200 0<br>A A G T A T A T A C T T T A T T C G | 880 |
| 11 | SRR7540368 | | | 121135..121152 | 80 60 80 60 80 60 0 40 80 60 40 0 0 0 0 40 200 0<br>A T G C A T T T A T C A A A T T C A | 880 |
| 12 | SRR7540368 | | | 142827..142844 | 80 60 0 40 80 60 80 60 80 60 40 0 0 0 0 40 200 0<br>A T C T A T A C A T C T C G A T C G | 880 |
| 13 | SRR7540368 | | | complement(99201..99218) | 80 60 0 40 80 60 80 60 80 60 40 0 0 0 0 40 200 0<br>A T C T A T A C A T C T C G A T C G | 880 |
| 14 | SRR7540368 | | | complement(80831..80848) | 80 40 80 60 80 60 0 40 80 60 60 0 0 0 0 40 200 0<br>A C G C A T C T A T T T G T G T C A | 880 |
| 15 | SRR7540368 | | | complement(73712..73729) | 80 60 80 40 80 40 80 0 80 60 40 0 0 0 0 40 200 0<br>A T G T A C A A A T C C A A A T C G | 880 |
| 16 | SRR7540368 | | | complement(49441..49458) | 80 60 80 40 80 60 80 40 80 40 0 0 0 0 0 40 200 0<br>A T G T A T A T A C G T T T A T C C | 880 |
| 17 | SRR7540368 | | | complement(39768..39785) | 80 60 80 0 80 60 80 60 80 0 60 0 0 0 0 40 200 0<br>A T G G A T A C A A T C C G C T C A | 880 |
| 18 | SRR7540368 | | | 80279..80296 | 80 60 80 40 80 60 80 40 80 60 0 0 0 0 0 15 200 0<br>A T G T A T A T A T A T T G G A C A | 875 |
| 19 | SRR7540368 | | | 121113..121130 | 80 40 80 60 80 40 80 60 80 60 0 0 0 0 0 15 200 0<br>A C G C A C A C A T A C T T C A C A | 875 |
| 20 | SRR7540368 | | | 79396..79413 | 80 60 80 0 80 60 0 60 80 60 60 0 0 0 0 40 200 0<br>A T G G A T T C A T T T G G T T C T | 860 |
| 21 | SRR7540368 | | | complement(121291..121308) | 80 60 0 60 80 60 80 0 80 60 60 0 0 0 0 40 200 0<br>A T T C A T A A A T T A T G G T C A | 860 |
| 22 | SRR7540368 | | | complement(70632..70649) | 80 60 80 0 80 40 80 60 20 60 60 0 0 0 0 40 200 0<br>A T G A A C A C G T T C A A T T C A | 860 |
| 23 | SRR7540368 | | | complement(61622..61639) | 80 60 0 40 80 60 80 40 80 60 40 0 0 0 0 40 200 0<br>A T C T A T A T A T C T A A A T C T | 860 |
| 24 | SRR7540368 | | | complement(60844..60861) | 80 60 80 40 80 0 20 60 80 60 60 0 0 0 0 40 200 0<br>A T G T A G G C A T T G G G T T C G | 860 |
| 25 | SRR7540368 | | | complement(31671..31688) | 80 60 80 60 80 60 20 40 80 0 60 0 0 0 0 40 200 0<br>A T G C A T G T A A T G G A A T C G | 860 |
| 26 | SRR7540368 | | | complement(2328..2345) | 80 60 80 60 80 0 80 0 80 60 40 0 0 0 0 40 200 0<br>A T G C A G A A A T C T T T T T C A | 860 |
| 27 | SRR7540368 | | | 6063..6080 | 80 60 80 40 80 40 80 0 80 60 40 0 0 0 0 15 200 0<br>A T G T A C A A A T C C A C A A C G | 855 |
| 28 | SRR7540368 | | | complement(83125..83142) | 80 60 80 60 0 40 80 40 80 60 60 0 0 0 0 15 200 0<br>A T G C T C A T A T T G G T G A C G | 855 |
| 29 | SRR7540368 | | | complement(81666..81683) | 80 60 0 60 80 60 80 40 80 60 40 0 0 0 0 15 200 0<br>A T C C A T A T A T C T C T G A C T | 855 |
| 30 | SRR7540368 | | | 97295..97312 | 80 0 80 60 80 60 80 60 20 60 60 0 0 0 0 5 200 0<br>A A G C A T A C G T T T C A T G C T | 845 |
| 31 | SRR7540368 | | | complement(144733..144750) | 80 0 80 60 80 60 80 60 20 60 60 0 0 0 0 5 200 0<br>A A G C A T A C G T T T C A T G C T | 845 |
| 32 | SRR7540368 | | | 1643..1660 | 80 40 0 40 80 60 80 40 80 40 60 0 0 0 0 40 200 0<br>A C T T A T A T A C T C G T G T C A | 840 |
| 33 | SRR7540368 | | | 14418..14435 | 80 40 20 40 80 60 80 60 80 60 0 0 0 0 0 40 200 0<br>A C A T A T A C A T G T C T T T C T | 840 |
| 34 | SRR7540368 | | | 19763..19780 | 20 40 80 60 80 60 0 60 80 60 60 0 0 0 0 40 200 0<br>G C G C A T T C A T T T G A T C C T | 840 |
| 35 | SRR7540368 | | | 33102..33119 | 80 60 20 40 80 0 80 40 80 60 60 0 0 0 0 40 200 0<br>A T A T A G A T A T T T T A T C C A | 840 |
| 36 | SRR7540368 | | | 55997..56014 | 80 60 80 40 80 0 80 0 80 60 40 0 0 0 0 40 200 0<br>A T G T A A A G A T C A T T T C C T | 840 |
| 37 | SRR7540368 | | | 68490..68507 | 80 60 80 0 80 60 0 40 80 60 60 0 0 0 0 40 200 0<br>A T G A A T C T A T T T T A T T C T | 840 |
| 38 | SRR7540368 | | | complement(121638..121655) | 0 40 80 40 80 40 80 40 80 60 60 0 0 0 0 40 200 0<br>C C G T A C A T A T T T G T T C C T | 840 |
| 39 | SRR7540368 | | | complement(56227..56244) | 80 60 0 60 80 60 80 0 80 60 40 0 0 0 0 40 200 0<br>A T T C A T A G A T C T G C G C C C | 840 |
| 40 | SRR7540368 | | | complement(46735..46752) | 80 60 80 0 80 60 80 60 0 40 60 0 0 0 0 40 200 0<br>A T G G A T A C T C T T A A A T C A | 840 |

**Back to input page**

CCTAGAGAGCCAGCATGTACA
Version: 3.12.0 using NCBI BLAST 2.2.26+
Lenz *et al.* 2010; Lenz and Knoop 2013; Lenz *et al.* 2018
Bug reports, feature requests and any comments go to hlenz (at) prepact.de
Imprint Privacy

**Fig. S3.** Weight matrix used for prediction of dPPRe$^{rpl2}$ cytidine targets within the *N. benthamiana* chloroplast genome by the TargetScan tool of PREPACT (Lenz et al., 2018) and the scanning output with the 40 best predicted targets. (*Top*) Each column corresponds to one nucleotide position and is labelled either (i) with type and fifth and last amino acid of the corresponding PPR or (ii) with the position relative to the editing site (EdS). Nucleotide distributions were set according to the published PPR-RNA binding code. Positional weighs are shown below. (*Bottom*) Best 40 predicted targets with the position in the genome and the matrix score (maximum 1000). Matches are colored in green and mismatches in red and given are the percentages of the match.

### DNA sequence of *dPPRe*$^{rpl2}$

**ATGGATTCTCAACTTGTTCTTTCTCTTAAGCTTAACCCTTCTTTTACTCCTCTTTCTCCTCTTTTTCCTTTTACTCCTTGTTC**
**TTCTTTTTCTCCTTCTCTTAGATTTTCTTCTTGTTATTCTAGGAGACTTTACTCTCCTGTTACTGTTTATGCTGCTAAGAAGC**
**TTTCTCATAAGATTTCTTCTGAGTTTGATGATAGAATC**GGTCTTATTAATATCTTCTACCCTTATGATGTGCCTGATTACGCT
GGATATCCTTATGATGTTCCTGATTATGCTGGATCTTATCCATATGATGTTCCAGATTATGCTGCTCAGTGTTCTAGTCTGCC
ATTGGATTCTCTATTGCTGCATTTGACAGCGCCGGCTCCTGCACCTGCACCGGCACCAAGACGATCCCATCAGACACCAACCC
CACCTCACTCTTTCCTGTCTCCAGACGCTCAGGTGTTGGTCCTTGCAATAAGCTCTCACCCGCTTCCGACACTTGCGGCATTC
TTGGCTTCTAGGCGTGATGAACTATTGAGAGCTGACATTACATCATTGCTTAAGGCCTTGGAATTGTCTGGCCACTGGGAATG
GGCGTTGGCGTTACTCAGATGGGCGGGAAAGGAAGGAGCTGCTGATGCATCCGCATTGGAAATGGTGGTCAGAGCTTTGGGAA
GGGAGGGTCAACATGATGCTGTGTGTGCTCTCCTTGATGAAACTCCTCTTCCTCCTGGGTCTCGTCTTGATGTTCGTGCATAT
ACGACAGTTTTACATGCTCTGTCGAGGGCGGGAAGGTACGAGCGAGCCTTAGAATTGTTCGCCGAATTAAGGCGTCAGGGAGT
GGCTCCTACTGTTGTTACGTATTCAACCTTGATCGATGGACTGTGTAAGGCCGGGAAGCTAGATGAAGCTCTTAAATTATTCG
AAGAGATGGTGGAGAAGGGTATCAAGCCAAATGTCGTAACATATAATACCCTAATCGATGGTCTGTGCAAGGCAGGTAAACTC
GATGAAGCCCTAAAATTGTTTGAGGAAATGGTAGAGAAAGGTATTAAACCGGATGTGGTGACGTACACGACTCTCATCGATGG
TCTTTGCAAAGCTGGGAAGTTGGACGAGGCTCTGAAATTGTTTGAAGAGATGGTAGAAAAAGGCATAAAACCAGATGTGGTCA
CTTATAACACACTTATAGACGGCTTGTGTAAAGCTGGGAAGTTGGACGAAGCTCTAAAGCTATTCGAGGAAATGGTTGAGAAA
GGAATTAAGCCATCAGTGGTTACCTACTCAACGCTGATCGATGGCCTATGTAAAGCCGGGAAACTTGATGAAGCACTTAAACT
CTTCGAGGAGATGGTGGAAAAAGGAATCAAGCCCAACGTTGTCACATACAATACGCTAATTGACGGCCTCTGTAAGGCGGGAA
AGCTGGATGAGGCTTTAAAACTTTTCGAAGAGATGGTGGAGAAAGGAATCAAACCGGATGTTGTAACGTATTCAACTTTAATC
GACGGGCTATGCAAGGCTGGGAAGCTCGATGAGGCCTTGAAGTTATTTGAAGAAATGGTGGAAAAAGGGATCAAACCAAATGT
GGTTACTTACAATACTCTCATCGATGGTCTCTGTAAGGCAGGGAAACTTGATGAGGCACTAAAGCTCTTCGAAGAGATGGTGG
AGAAAGGTATAAAGCCTTCAGTAGTTACGTACTCAACGCTGATCGACGGCTTGGCAAAGTGTGGAAAACTCGACGAGGCACTT
AAACTCTTTGAAGAGATGGTCGAAAAGGGGATAAAGCCTAATGTCGTTACATATAATACGCTAATCGACGGCTTGTGCAAAGC
GGGAAAACTAGATGAGGCTTTGAAACTCTTTGAGGAGATGGTTGAGAAGGGAATCAAGCCGGACGTGGTTTCATGGAATGCTA
TGATCTCTGGATACGCAATGCATGGACATGGAAAGGAAGCACTCGAGCTATTTGAAGAGATGCAGCAGTCTGGAATCAAGCCT
GATCATGTTACGTTTACTGGAGTCCTATCTGCTTGCTCTCATGCTGGACTCGTTGACGAAGGAAGGCAGTACTTTAATTCAAT
GAAGAAAGACTACGGAATCGAGCCTAGGGTTGAACATTACGGCTGCATGGTTGACCTGCTTGGAAGGGCTGGAAGGCTAGACG
AGGCATACGAATTTATCGAGTCAATGCCTATCGAACCTAACGAAGCAACTTGGGGAGCCCTGCTTGGTTCTTGTAGAACTTAT
GGTAACGTAGAGTTGGGCGAACTTGTTGCGAAAGAACGTCTGAAATTGGATCCGAAAAATGCTGCTACCTATGTTCTCCTTTC
CAACATCTATGCTGAAGCTGGAAAGTGGGACATGGTATCATGGGTGCGGACTATGATGCGCGAGAGAGGGATTCGCAAGGAGC
CAGGACGTAGCTGGATCGAAGTCGATAACAAGATTCATGATTTTTTGGTAGCCGACTCATCTCATCCAGAATGCAAGGAGATA
AATGAAAGTAAGGACAAAGTGATTGAAAAAATAAAGGCTGAAGGGTACATTCCTGATACCCGGCTAGTGCTGAAGAACAAGAA
CATGAAGGACAAAGAACTTGATATTTGCTCTCACAGTGAGAAGTTGGCAATTGTGTATGGGCTGATGCATACACCCCCTGGAA
ATCCTATTCGTGTATTCAAGAACCTTCGGGTGTGCACCGATTGTCATGGAGCAACTAAGTTAATCTCAAAGGTGGAGGGACGA
GAGATCATAGTGAGGGATGCTAATCGGTTTCACCATTTTAAAGATGGCGTCTGCTCATGTGGTGATTACTGGTAG

### Amino acid sequence of dPPRe$^{rpl2}$

**MDSQLVLSLKLNPSFTPLSPLFPFTPCSSFSPSLRFSSCYSRRLYSPVTVYAAKKLSHKISSEFDDRI**GLINIFYPYDVPDYA
GYPYDVPDYAGSYPYDVPDYAAQCSSLPLDSLLLHLTAPAPAPAPAPRRSHQTPTPPHSFLSPDAQVLVLAISSHPLPTLAAF
LASRRDELLRADITSLLKALELSGHWEWALALLRWAGKEGAADASALEMVVRALGREGQHDAVCALLDETPLPPGSRLDVRAY
TTVLHALSRAGRYERALELFAELRRQGVAPTVVTYSTLIDGLCKAGKLDEALKLFEEMVEKGIKPNVVTYNTLIDGLCKAGKL
DEALKLFEEMVEKGIKPDVVTYTTLIDGLCKAGKLDEALKLFEEMVEKGIKPDVVTYNTLIDGLCKAGKLDEALKLFEEMVEK
GIKPSVVTYSTLIDGLCKAGKLDEALKLFEEMVEKGIKPNVVTYNTLIDGLCKAGKLDEALKLFEEMVEKGIKPDVVTYSTLI
DGLCKAGKLDEALKLFEEMVEKGIKPNVVTYNTLIDGLCKAGKLDEALKLFEEMVEKGIKPSVVTYSTLIDGLAKCGKLDEAL
KLFEEMVEKGIKPNVVTYNTLIDGLCKAGKLDEALKLFEEMVEKGIKPDVVSWNAMISGYAMHGHGKEALELFEEMQQSGIKP
DHVTFTGVLSACSHAGLVDEGRQYFNSMKKDYGIEPRVEHYGCMVDLLGRAGRLDEAYEFIESMPIEPNEATWGALLGSCRTY
GNVELGELVAKERLKLDPKNAATYVLLSNIYAEAGKWDMVSWVRTMMRERGIRKEPGRSWIEVDNKIHDFLVADSSHPECKEI
NESKDKVIEKIKAEGYIPDTRLVLKNKNMKDKELDICSHSEKLAIVYGLMHTPPGNPIRVFKNLRVCTDCHGATKLISKVEGR
EIIVRDANRFHHFKDGVCSCGDYW

### DNA sequence of *dPPRe*$^{ndhB}$

**ATGGATTCTCAACTTGTTCTTTCTCTTAAGCTTAACCCTTCTTTTACTCCTCTTTCTCCTCTTTTTCCTTTTACTCCTTGTTC**
**TTCTTTTTCTCCTTCTCTTAGATTTTCTTCTTGTTATTCTAGGAGACTTTACTCTCCTGTTACTGTTTATGCTGCTAAGAAGC**
**TTTCTCATAAGATTTCTTCTGAGTTTGATGATAGAATC**GGTCTTATTAATATCTTCTACCCTTATGATGTGCCTGATTACGCT
GGATATCCTTATGATGTTCCTGATTATGCTGGATCTTATCCATATGATGTTCCAGATTATGCTGCTCAGTGTTCTAGTCTGCC
ATTGGATTCTCTATTGCTGCATTTGACAGCGCCGGCTCCTGCACCTGCACCGGCACCAAGACGATCCCATCAGACACCAACCC
CACCTCACTCTTTCCTGTCTCCAGACGCTCAGGTGTTGGTCCTTGCAATAAGCTCTCACCCGCTTCCGACACTTGCGGCATTC
TTGGCTTCTAGGCGTGATGAACTATTGAGAGCTGACATTACATCATTGCTTAAGGCCTTGGAATTGTCTGGCCACTGGGAATG
GGCGTTGGCGTTACTCAGATGGGCGGGAAAGGAAGGAGCTGCTGATGCATCCGCATTGGAAATGGTGGTCAGAGCTTTGGGAA
GGGAGGGTCAACATGATGCTGTGTGTGCTCTCCTTGATGAAACTCCTCTTCCTCCTGGGTCTCGTCTTGATGTTCGTGCATAT
ACGACAGTTTTACATGCTCTGTCGAGGGCGGGAAGGTACGAGCGAGCCTTAGAATTGTTCGCCGAATTAAGGCGTCAGGGAGT
GGCTCCTACTGTTGTTACGTATAATACCTTGATCGATGGACTGTGTAAGGCCGGGAAGCTAGATGAAGCTCTTAAATTATTCG
AAGAGATGGTGGAGAAGGGTATCAAGCCAGATGTCGTAACATATAATACCCTAATCGATGGTCTGTGCAAGGCAGGTAAACTC
GATGAAGCCCTAAAATTGTTTGAGGAAATGGTAGAGAAAGGTATTAAACCGTCGGTGGTGACGTACAATACTCTCATCGATGG
TCTTTGCAAAGCTGGGAAGTTGGACGAGGCTCTGAAATTGTTTGAAGAGATGGTAGAAAAAGGCATAAAACCAGATGTGGTCA
CTTATAACACACTTATAGACGGCTTGTGTAAAGCTGGGAAGTTGGACGAAGCTCTAAAGCTATTCGAGGAAATGGTTGAGAAA
GGAATTAAGCCATCAGTGGTTACCTACTCAACGCTGATCGATGGCCTATGTAAAGCCGGGAAACTTGATGAAGCACTTAAACT

CTTCGAGGAGATGGTGGAAAAAGGAATCAAGCCCAACGTTGTCACATACAATACGCTAATTGACGGCCTCTGTAAGGCGGGAA
AGCTGGATGAGGCTTTAAAACTTTTCGAAGAGATGGTGGAGAAAGGAATCAAACCGGATGTTGTAACGTATAATACTTTAATC
GACGGGCTATGCAAGGCTGGGAAGCTCGATGAGGCCTTGAAGTTATTTGAAGAAATGGTGGAAAAAGGGATCAAACCAGATGT
GGTTACTTACACGACTCTCATCGATGGTCTCTGTAAGGCAGGGAAACTTGATGAGGCACTAAAGCTCTTCGAAGAGATGGTGG
AGAAAGGTATAAAGCCTGATGTAGTTACGTACAATACGCTGATCGACGGCTTGGCAAAGTGTGGAAAACTCGACGAGGCACTT
AAACTCTTTGAAGAGATGGTCGAAAAGGGGATAAAGCCTTCAGTCGTTACATATAATACGCTAATCGACGGCTTGTGCAAAGC
GGGAAAACTAGATGAGGCTTTGAAACTCTTTGAGGAGATGGTTGAGAAGGGAATCAAGCCGGACGTGGTTTCATGGTCAGCTA
TGATCTCTGGATACGCAATGCATGGACATGGAAAGGAAGCACTCGAGCTATTTGAAGAGATGCAGCAGTCTGGAATCAAGCCT
AACCATGTTACGTTTACTGGAGTCCTATCTGCTTGCTCTCATGCTGGACTCGTTGACGAAGGAAGGCAGTACTTTAATTCAAT
GAAGAAAGACTACGGAATCGAGCCTAGGGTTGAACATTACGGCTGCATGGTTGACCTGCTTGGAAGGGCTGGAAGGCTAGACG
AGGCATACGAATTTATCGAGTCAATGCCTATCGAACCTAACGAAGCAACTTGGGGAGCCCTGCTTGGTTCTTGTAGAACTTAT
GGTAACGTAGAGTTGGGCGAACTTGTTGCGAAAGAACGTCTGAAATTGGATCCGAAAAATGCTGCTACCTATGTTCTCCTTTC
CAACATCTATGCTGAAGCTGGAAAGTGGGACATGGTATCATGGGTGCGGACTATGATGCGCGAGAGAGGGATTCGCAAGGAGC
CAGGACGTAGCTGGATCGAAGTCGATAACAAGATTCATGATTTTTTGGTAGCCGACTCATCTCATCCAGAATGCAAGGAGATA
AATGAAAGTAAGGACAAAGTGATTGAAAAAATAAAGGCTGAAGGGTACATTCCTGATACCCGGCTAGTGCTGAAGAACAAGAA
CATGAAGGACAAAGAACTTGATATTTGCTCTCACAGTGAGAAGTTGGCAATTGTGTATGGGCTGATGCATACACCCCCTGGAA
ATCCTATTCGTGTATTCAAGAACCTTCGGGTGTGCACCGATTGTCATGGAGCAACTAAGTTAATCTCAAAGGTGGAGGGACGA
GAGATCATAGTGAGGGATGCTAATCGGTTTCACCATTTTAAAGATGGCGTCTGCTCATGTGGTGATTACTGGTAG

### Amino acid sequence of dPPRe$^{ndhB}$

**MDSQLVLSLKLNPSFTPLSPLFPFTPCSSFSPSLRFSSCYSRRLYSPVTVYAAKKLSHKISSEFDDRI**GLINIFYPYDVPDYA
GYPYDVPDYAGSYPYDVPDYAAQCSSLPLDSLLLHLTAPAPAPAPAPRRSHQTPTPPHSFLSPDAQVLVLAISSHPLPTLAAF
LASRRDELLRADITSLLKALELSGHWEWALALLRWAGKEGAADASALEMVVRALGREGQHDAVCALLDETPLPPGSRLDVRAY
TTVLHALSRAGRYERALELFAELRRQGVAPTVVTYNTLIDGLCKAGKLDEALKLFEEMVEKGIKPDVVTYNTLIDGLCKAGKL
DEALKLFEEMVEKGIKPSVVTYNTLIDGLCKAGKLDEALKLFEEMVEKGIKPDVVTYNTLIDGLCKAGKLDEALKLFEEMVEK
GIKPSVVTYSTLIDGLCKAGKLDEALKLFEEMVEKGIKPNVVTYNTLIDGLCKAGKLDEALKLFEEMVEKGIKPDVVTYNTLI
DGLCKAGKLDEALKLFEEMVEKGIKPDVVTYTTLIDGLCKAGKLDEALKLFEEMVEKGIKPDVVTYNTLIDGLAKCGKLDEAL
KLFEEMVEKGIKPSVVTYNTLIDGLCKAGKLDEALKLFEEMVEKGIKPDVVSWSAMISGYAMHGHGKEALELFEEMQQSGIKP
NHVTFTGVLSACSHAGLVDEGRQYFNSMKKDYGIEPRVEHYGCMVDLLGRAGRLDEAYEFIESMPIEPNEATWGALLGSCRTY
GNVELGELVAKERLKLDPKNAATYVLLSNIYAEAGKWDMVSWVRTMMRERGIRKEPGRSWIEVDNKIHDFLVADSSHPECKEI
NESKDKVIEKIKAEGYIPDTRLVLKNKNMKDKELDICSHSEKLAIVYGLMHTPPGNPIRVFKNLRVCTDCHGATKLISKVEGR
EIIVRDANRFHHFKDGVCSCGDYW

### DNA sequence of *dPPRe$^{nad7}$*

**ATGTTGTCTCTTAGACAGTCTATTAGATTTTTCAAGCCTGCTACCAGAACTTTGTGCTCTTCTAGGTATCTTCTTCAACAAAA**
**GCCT**GGTCTTATTAATATCTTCTACCCTTATGATGTGCCTGATTACGCTGGATATCCTTATGATGTTCCTGATTATGCTGGAT
CTTATCCATATGATGTTCCAGATTATGCTGCTCAGTGTTCTAGTCTGCCATTGGATTCTCTATTGCTGCATTTGACAGCGCCG
GCTCCTGCACCTGCACCGGCACCAAGACGATCCCATCAGACACCAACCCCACCTCACTCTTTCCTGTCTCCAGACGCTCAGGT
GTTGGTCCTTGCAATAAGCTCTCACCCGCTTCCGACACTTGCGGCATTCTTGGCTTCTAGGCGTGATGAACTATTGAGAGCTG
ACATTACATCATTGCTTAAGGCCTTGGAATTGTCTGGCCACTGGGAATGGGCGTTGGCGTTACTCAGATGGGCGGGAAAGGAA
GGAGCTGCTGATGCATCCGCATTGGAAATGGTGGTCAGAGCTTTGGGAAGGGAGGGTCAACATGATGCTGTGTGTGCTCTCCT
TGATGAAACTCCTCTTCCTCCTGGGTCTCGTCTTGATGTTCGTGCATATACGACAGTTTTACATGCTCTGTCGAGGGCGGGAA
GGTACGAGCGAGCCTTAGAATTGTTCGCCGAATTAAGGCGTCAGGGAGTGGCTCCTACTGTTGTTACGTATACTACCTTGATC
GATGGACTGTGTAAGGCCGGGAAGCTAGATGAAGCTCTTAAATTATTCGAAGAGATGGTGGAGAAGGGTATCAAGCCAGACGT
CGTAACATATAATACCCTAATCGATGGTCTGTGCAAGGCAGGTAAACTCGATGAAGCCCTAAAATTGTTTGAGGAAATGGTAG
AGAAAGGTATTAAACCGGATGTGGTGACGTACTCGACTCTCATCGATGGTCTTTGCAAAGCTGGGAAGTTGGACGAGGCTCTG
AAATTGTTTGAAGAGATGGTAGAAAAAGGCATAAAACCAAATGTGGTCACTTATAACACACTTATAGACGGCTTGTGTAAAGC
TGGGAAGTTGGACGAAGCTCTAAAGCTATTCGAGGAAATGGTTGAGAAAGGAATTAAGCCATCAGTGGTTACCTACTCAACGC
TGATCGATGGCCTATGTAAAGCCGGGAAACTTGATGAAGCACTTAAACTCTTCGAGGAGATGGTGGAAAAAGGAATCAAGCCC
AACGTTGTCACATACTCGACGCTAATTGACGGCCTCTGTAAGGCGGGAAAGCTGGATGAGGCTTTAAAACTTTTCGAAGAGAT
GGTGGAGAAAGGAATCAAACCGAATGTTGTAACGTATTCAACTTTAATCGACGGGCTATGCAAGGCTGGGAAGCTCGATGAGG
CCTTGAAGTTATTTGAAGAAATGGTGGAAAAAGGGATCAAACCAAATGTGGTTACTTACAGCACTCTCATCGATGGTCTCTGT
AAGGCAGGGAAACTTGATGAGGCACTAAAGCTCTTCGAAGAGATGGTGGAGAAAGGTATAAAGCCTAACGTAGTTACGTACAA
CACGCTGATCGACGGCTTGGCAAAGTGTGGAAAACTCGACGAGGCACTTAAACTCTTTGAAGAGATGGTCGAAAAGGGGATAA
AGCCTAGCGTCGTTACATATAATACGCTAATCGACGGCTTGTGCAAAGCGGGAAAACTAGATGAGGCTTTGAAACTCTTTGAG
GAGATGGTTGAGAAGGGAATCAAGCCGGACGTGGTTTCATGGAATGCTATGATCTCTGGATACGCAATGCATGGACATGGAAA
GGAAGCACTCGAGCTATTTGAAGAGATGCAGCAGTCTGGAATCAAGCCTGATCATGTTACGTTTACTGGAGTCCTATCTGCTT
GCTCTCATGCTGGACTCGTTGACGAAGGAAGGCAGTACTTTAATTCAATGAAGAAAGACTACGGAATCGAGCCTAGGGTTGAA
CATTACAATTGCATGGTTGACCTGCTTGGAAGGGCTGGAAGGCTAGACGAGGCATACGAATTTATCGAGTCAATGCCTATCGA
ACCTGATGAAGCAACTTGGGGAGCCCTGCTTGGTTCTTGTAGAACTTATGGTAACGTAGAGTTGGGCGAACTTGTTGCGAAAG
AACGTCTGAAATTGGATCCGAAAAATGCTGCTACCTATGTTCTCCTTTCCAACATCTATGCTGAAGCTGGAAAGTGGGACATG
GTATCATGGGTGCGGACTATGATGCGCGAGAGAGGGATTCGCAAGGAGCCAGGACGTAGCTGGATCGAAGTCGATAACAAGAT
TCATGATTTTTTGGTAGCCGACTCATCTCATCCAGAATGCAAGGAGATAAATGAAAGTAAGGACAAAGTGATTGAAAAAATAA
AGGCTGAAGGGTACATTCCTGATACCCGGCTAGTGCTGAAGAACAAGAACATGAAGGACAAAGAACTTGATATTTGCTCTCAC
AGTGAGAAGTTGGCAATTGTGTATGGGCTGATGCATACACCCCCTGGAAATCCTATTCGTGTATTCAAGAACCTTCGGGTGTG
CACCGATTGTCATGGAGCAACTAAGTTAATCTCAAAGGTGGAGGGACGAGAGATCATAGTGAGGGATGCTAATCGGTTTCACC
ATTTTAAAGATGGCGTCTGCTCATGTGGTGATTACTGGTAG

**Amino acid sequence of dPPRe*nad7***

**MLSLRQSIRFFKPATRTLCSSRYLLQQKP**GLINIFYPYDVPDYAGYPYDVPDYAGSYPYDVPDYAAQCSSLPLDSLLLHLTAP
APAPAPAPRRSHQTPTPPHSFLSPDAQVLVLAISSHPLPTLAAFLASRRDELLRADITSLLKALELSGHWEWALALLRWAGKE
GAADASALEMVVRALGREGQHDAVCALLDETPLPPGSRLDVRAYTTVLHALSRAGRYERALELFAELRRQGVAPTVVTYTTLI
DGLCKAGKLDEALKLFEEMVEKGIKPDVVTYNTLIDGLCKAGKLDEALKLFEEMVEKGIKPDVVTYSTLIDGLCKAGKLDEAL
KLFEEMVEKGIKPNVVTYNTLIDGLCKAGKLDEALKLFEEMVEKGIKPSVVTYSTLIDGLCKAGKLDEALKLFEEMVEKGIKP
NVVTYSTLIDGLCKAGKLDEALKLFEEMVEKGIKPNVVTYSTLIDGLCKAGKLDEALKLFEEMVEKGIKPNVVTYSTLIDGLC
KAGKLDEALKLFEEMVEKGIKPNVVTYNTLIDGLAKCGKLDEALKLFEEMVEKGIKPSVVTYNTLIDGLCKAGKLDEALKLFE
EMVEKGIKPDVVSWNAMISGYAMHGHGKEALELFEEMQQSGIKPDHVTFTGVLSACSHAGLVDEGRQYFNSMKKDYGIEPRVE
HYNCMVDLLGRAGRLDEAYEFIESMPIEPDEATWGALLGSCRTYGNVELGELVAKERLKLDPKNAATYVLLSNIYAEAGKWDM
VSWVRTMMRERGIRKEPGRSWIEVDNKIHDFLVADSSHPECKEINESKDKVIEKIKAEGYIPDTRLVLKNKNMKDKELDICSH
SEKLAIVYGLMHTPPGNPIRVFKNLRVCTDCHGATKLISKVEGREIIVRDANRFHHFKDGVCSCGDYW

**Fig. S4.** Sequences of dPPRe*rpl2*, dPPRe*ndhB* and dPPRe*nad7*. Arabidopsis RecA and yeast Cox4 transit peptides are in bold black, the 3xHA tag is shown in red and the synthetic P-PPR array is delimited by a box. The C-terminal synthetic P2, S2, L2 and PPR56's E1-E2-DYW are highlighted in turquoise, pink, yellow and grey, respectively. The starting amino acid of the dPPR protein expressed as a recombinant protein in *E. coli* is shaded in red.

| | rep1 | rep2 | rep3 | Mean | SD |
|---|---|---|---|---|---|
| **rdPPRe$^{rpl2}$** | 0,62 | 0,59 | 0,49 | 0,566667 | 0,068069 |
| **rdPPRe$^{ndhB}$** | 0,3 | 0,37 | 0,31 | 0,326667 | 0,037859 |

**Table S1.** dPPRe editing efficiency in *E. coli* (Sanger).

| **dPPRe$^{rpl2}$ 2dpi** | **Editing rate** | **P19 (control) 2dpi** | **Editing rate** | **Unpaired *t* test results**<br>**dPPRe$^{rpl2}$ 2dpi vs P19** |
|---|---|---|---|---|
| **rep 1** | 0,45361991 | **rep 1** | 0 | two-tailed P value = 0.0006 |
| **rep 2** | 0,368159204 | **rep 2** | 0 | t = 9.7696 |
| **rep 3** | 0,32038835 | **rep 3** | 0 | df = 4 |
| **Mean** | **0,380722488** | **Mean** | **0** | standard error of difference = 0.039 |
| **SD** | **0,067498439** | **SD** | **0** | |

| **dPPRe$^{rpl2}$ 3dpi** | **Editing rate** | **P19 (control) 3dpi** | **Editing rate** | **dPPRe$^{rpl2}$ 3dpi vs P19** |
|---|---|---|---|---|
| **rep 1** | 0,488207547 | **rep 1** | 0 | two-tailed P value < 0.0001 |
| **rep 2** | 0,61026616 | **rep 2** | 0 | t = 12.8194 |
| **rep 3** | 0,6918429 | **rep 3** | 0 | df = 6 |
| **rep 4** | 0,528205128 | **rep 4** | 0 | standard error of difference = 0.045 |
| **Mean** | **0,579630434** | **Mean** | 0 | |
| **SD** | **0,090430252** | **SD** | 0 | |

| **dPPRe$^{rpl2}$ 2dpi vs 3dpi** |
|---|
| two-tailed P value = 0.0247 |
| t = 3.1748 |
| df = 5 |
| standard error of difference = 0.063 |

| **dPPRe$^{ndhB}$ 3dpi** | **Editing rate** | **P19 (control) 3dpi** | **Editing rate** | **Unpaired *t* test results**<br>**dPPRe$^{ndhB}$ 3dpi vs P19** |
|---|---|---|---|---|
| **rep 1** | 0,230985915 | **rep 1** | 0 | two-tailed P value < 0.0001 |
| **rep 2** | 0,160206718 | **rep 2** | 0 | t = 9.4815 |
| **rep 3** | 0,206145967 | **rep 3** | 0 | df = 9 |
| **rep 4** | 0,12006079 | **rep 4** | 0 | standard error of difference = 0.018 |
| **rep 5** | 0,14619883 | **rep 5** | 0 | |
| **Mean** | **0,172719644** | **rep 6** | 0 | |
| **SD** | **0,045125268** | **Mean** | **0** | |
| | | **SD** | **0** | |

| **dPPRe$^{nad7}$ 3dpi** | **Editing rate** | **P19 (control) 3dpi** | **Editing rate** | **Unpaired *t* test results**<br>**dPPRe$^{nad7}$ 3dpi vs P19** |
|---|---|---|---|---|
| **rep 1** | 0,314159292 | **rep 1** | 0 | two-tailed P value < 0.0001 |
| **rep 2** | 0,349112426 | **rep 2** | 0 | t = 11.2182 |
| **rep 3** | 0,493723849 | **rep 3** | 0 | df = 8 |
| **rep 4** | 0,369909502 | **rep 4** | 0 | standard error of difference = 0.036 |
| **rep 5** | 0,476629361 | **rep 5** | 0 | |
| **Mean** | **0,400706886** | **Mean** | **0** | |
| **SD** | **0,079870725** | **SD** | **0** | |

**Table S2.** dPPRe editing efficiency *in planta* (Sanger).

| | baseMean | log2FoldChange | lfcSE | stat | pvalue | padj |
|---|---|---|---|---|---|---|
| gene-blatn_trnI_1 | 2411,217493 | -0,214996618 | 0,208204434 | -1,0326227 | 0,301780516 | 0,653109223 |
| gene-blatn_trnL_1 | 668,6341773 | -0,185269079 | 0,232671599 | -0,7962686 | 0,425875956 | 0,694505405 |
| gene-blatn_trnV_1 | 585,1995695 | -0,118830208 | 0,207582182 | -0,572449 | 0,567017843 | 0,775821811 |
| gene-blatn_rrn23_1 | 17769,97961 | -0,316154653 | 0,33552088 | -0,9422801 | 0,346049247 | 0,658461475 |
| gene-blatn_rrn4.5_1 | 184,6078079 | -1,143964693 | 0,417976596 | -2,7369109 | 0,006201909 | 0,204960924 |
| gene-blatn_rrn5_1 | 51,62263707 | -0,38381381 | 0,379485638 | -1,0114054 | 0,311822455 | 0,653109223 |
| gene-blatn_trnR_1 | 6405,817524 | -0,634900099 | 0,257110254 | -2,469369 | 0,013535155 | 0,204960924 |
| gene-blatn_trnN_1 | 2018,865211 | -0,49291557 | 0,196281429 | -2,5112695 | 0,012029781 | 0,204960924 |
| gene-blatn_trnP_1 | 18281,21342 | -0,056034249 | 0,291375084 | -0,1923097 | 0,847499638 | 0,907423855 |
| gene-blatn_trnW_1 | 12661,49331 | 0,148720895 | 0,30082494 | 0,49437688 | 0,62104004 | 0,793135472 |
| gene-blatn_trnM_1 | 1929,800719 | 0,172424394 | 0,154998206 | 1,11242832 | 0,265954035 | 0,653109223 |
| gene-blatn_trnF_1 | 2193,697675 | -0,353571984 | 0,421582903 | -0,8386772 | 0,401650457 | 0,665233569 |
| gene-blatn_trnT_1 | 1443,108993 | -0,260708555 | 0,167013475 | -1,5610031 | 0,118523015 | 0,613582901 |
| gene-blatn_trnS_1 | 5994,060664 | 0,085666382 | 0,196078606 | 0,43689816 | 0,662185207 | 0,818244614 |
| gene-blatn_trnfM_1 | 504,4305075 | -0,590520778 | 0,226242058 | -2,6101282 | 0,009050829 | 0,204960924 |
| gene-blatn_trnG_1 | 1303,664561 | -0,403302752 | 0,229774729 | -1,7552093 | 0,079223506 | 0,559846109 |
| gene-blatn_trnE_1 | 6643,300004 | -0,384570582 | 0,259259103 | -1,4833446 | 0,137982888 | 0,613582901 |
| gene-blatn_trnY_1 | 15074,58935 | -0,324820777 | 0,214396779 | -1,515045 | 0,129760999 | 0,613582901 |
| gene-blatn_trnD_1 | 14348,98726 | -0,313625801 | 0,198272369 | -1,5817928 | 0,113696885 | 0,613582901 |
| gene-blatn_trnC_1 | 3295,593982 | -0,241991835 | 0,186343341 | -1,2986342 | 0,194069497 | 0,637866774 |
| gene-blatn_trnQ_1 | 1165,836718 | -0,141150295 | 0,225854617 | -0,6249609 | 0,531996747 | 0,762049394 |
| gene-blatn_trnH_1 | 1485,437541 | 0,069753196 | 0,174650295 | 0,39938779 | 0,689607487 | 0,820275417 |
| gene-blatn_trnA_1 | 182272,7798 | -0,209133802 | 0,239438024 | -0,8734361 | 0,382425449 | 0,658461475 |
| gene-blatn_trnK_1 | 490697,0434 | 0,467713872 | 0,198979431 | 2,35056393 | 0,018744987 | 0,248371083 |
| gene-blatn_rrn16_1 | 6077,72385 | -0,383645191 | 0,379401727 | -1,0111846 | 0,311928075 | 0,653109223 |
| gene-blatx_rpl23_1 | 10340,70468 | 0,250562422 | 0,213445863 | 1,17389215 | 0,240438229 | 0,637866774 |
| gene-blatx_ycf15_1 | 7565,42931 | -0,092592833 | 0,206667854 | -0,4480273 | 0,654133521 | 0,818244614 |
| gene-blatx_ndhE_1 | 66862,46382 | -0,13542248 | 0,209581371 | -0,646157 | 0,518177653 | 0,759627173 |
| gene-blatx_rpl32_1 | 25123,29377 | -0,25606553 | 0,217587425 | -1,1768397 | 0,239259475 | 0,637866774 |
| gene-blatx_rps19_1 | 5434,15878 | 0,542868965 | 0,30648568 | 1,77127024 | 0,076515773 | 0,559846109 |
| gene-blatx_rpl36_1 | 7769,164215 | 0,26678671 | 0,279693094 | 0,95385519 | 0,340156947 | 0,658461475 |
| gene-blatx_petD_1 | 276583,7855 | -0,116744081 | 0,205544307 | -0,5679753 | 0,57005177 | 0,775821811 |
| gene-blatx_psbH_1 | 125579,5681 | -0,028872397 | 0,181916298 | -0,1587125 | 0,873895359 | 0,921966437 |
| gene-blatx_psbN_1 | 43655,95947 | 0,121823054 | 0,162885261 | 0,74790716 | 0,454516156 | 0,708510479 |
| gene-blatx_psbT_1 | 65357,60068 | 0,029879207 | 0,137613504 | 0,21712409 | 0,828111641 | 0,895712591 |
| gene-blatx_clpP1_1 | 44224,4245 | 0,046544067 | 0,169787193 | 0,27413179 | 0,783983345 | 0,856724068 |
| gene-blatx_rpl20_1 | 8823,511236 | -0,015001924 | 0,150442548 | -0,0997186 | 0,920567714 | 0,947380366 |
| gene-blatx_rps18_1 | 30425,98101 | 0,288857149 | 0,142541396 | 2,02647903 | 0,04271572 | 0,411624208 |
| gene-blatx_rpl33_1 | 16984,78703 | 0,359193223 | 0,143563489 | 2,50198172 | 0,01235003 | 0,204960924 |
| gene-blatx_psaJ_1 | 126067,0736 | -0,192186146 | 0,271320375 | -0,7083366 | 0,478736265 | 0,724943488 |
| gene-blatx_petG_1 | 28224,51554 | -0,354817012 | 0,2737867 | -1,2959615 | 0,194988762 | 0,637866774 |
| gene-blatx_psbE_1 | 398339,2269 | -0,240642574 | 0,251996628 | -0,9549436 | 0,339606206 | 0,658461475 |
| gene-blatx_psbF_1 | 309251,4066 | -0,267320163 | 0,253072109 | -1,0563004 | 0,290831005 | 0,653109223 |
| gene-blatx_psbL_1 | 310270,2682 | -0,260634845 | 0,252922371 | -1,0304934 | 0,302778428 | 0,653109223 |
| gene-blatx_psbJ_1 | 197515,2287 | -0,230379329 | 0,248815193 | -0,9259054 | 0,354495148 | 0,658461475 |
| gene-blatx_psaI_1 | 71919,93762 | -0,250361482 | 0,251961391 | -0,9936502 | 0,320393204 | 0,653109223 |
| gene-blatx_ndhJ_1 | 32908,97888 | -0,1380485 | 0,352306142 | -0,3918424 | 0,695174632 | 0,820275417 |
| gene-blatx_rps14_1 | 96753,85243 | 0,103276543 | 0,206646314 | 0,49977442 | 0,617233921 | 0,793135472 |
| gene-blatx_psbZ_1 | 63117,06824 | -0,005269794 | 0,134971141 | -0,0390439 | 0,96885542 | 0,96885542 |
| gene-blatx_psbM_1 | 139983,0479 | -0,46520505 | 0,242878622 | -1,9153808 | 0,055443958 | 0,48975496 |
| gene-blatx_petN_1 | 83000,54267 | -0,184528298 | 0,325593421 | -0,5667446 | 0,570887747 | 0,775821811 |
| gene-blatx_atpH_1 | 122655,536 | 0,046247699 | 0,155935457 | 0,29658231 | 0,766785415 | 0,849937162 |
| gene-blatx_psbI_1 | 144781,0357 | -0,187645878 | 0,293879264 | -0,6385135 | 0,523139468 | 0,759627173 |
| gene-blatx_psbK_1 | 201091,4219 | -0,218738019 | 0,304160049 | -0,7191543 | 0,47204583 | 0,724943488 |
| gene-blatx_ndhB_1 | 114214,3108 | 0,197400415 | 0,17086314 | 1,15531304 | 0,247962257 | 0,641073152 |
| gene-blatx_rpl2_1 | 39848,53454 | 0,320652537 | 0,235931395 | 1,35909227 | 0,174117351 | 0,637866774 |
| gene-blatx_atpF_1 | 194547,2435 | 0,143364408 | 0,134263233 | 1,06778606 | 0,28561703 | 0,653109223 |
| gene-blatx_ycf3_1 | 90012,51427 | 0,023412992 | 0,169469637 | 0,1381545 | 0,890118317 | 0,925024917 |
| gene-blatx_psbB_1 | 377851,1366 | -0,045507897 | 0,150411682 | -0,3025556 | 0,762228552 | 0,849937162 |

| | | | | | | |
|---|---|---|---|---|---|---|
| gene-blatx_ycf2_1 | 46978,62049 | 0,025541666 | 0,325586255 | 0,07844823 | 0,937471508 | 0,955499807 |
| gene-blatx_petA_1 | 56436,16773 | 0,106829743 | 0,192577957 | 0,55473505 | 0,579075897 | 0,776987912 |
| gene-blatx_atpA_1 | 173994,2908 | 0,359013676 | 0,237167101 | 1,51375834 | 0,130087143 | 0,613582901 |
| gene-blatx_psaA_1 | 349664,9537 | 0,283714651 | 0,227066617 | 1,2494776 | 0,211490442 | 0,637866774 |
| gene-blatx_ndhK_1 | 39838,05547 | -0,188063557 | 0,37677688 | -0,4991377 | 0,617682357 | 0,793135472 |
| gene-blatx_atpI_1 | 76656,34709 | 0,074273071 | 0,170903114 | 0,43459168 | 0,663858838 | 0,818244614 |
| gene-blatx_psaB_1 | 367506,4078 | 0,248436456 | 0,218491923 | 1,13705099 | 0,255516972 | 0,644876168 |
| gene-blatx_rps2_1 | 7737,329305 | -0,057388895 | 0,196069864 | -0,2926962 | 0,76975441 | 0,849937162 |
| gene-blatx_rpoB_1 | 15685,25912 | 0,195979969 | 0,259621546 | 0,75486789 | 0,450328253 | 0,708510479 |
| gene-blatx_rps4_1 | 26861,10381 | 0,027014423 | 0,176680797 | 0,1528996 | 0,878477454 | 0,921966437 |
| gene-blatx_ndhH_1 | 18198,75448 | 0,422711965 | 0,26364618 | 1,60333052 | 0,108861703 | 0,613582901 |
| gene-blatx_ycf4_1 | 44547,48642 | 0,279837658 | 0,216904511 | 1,29014218 | 0,197001297 | 0,637866774 |
| gene-blatx_psbD_1 | 474468,1043 | -0,110607547 | 0,167335394 | -0,6609931 | 0,508616716 | 0,759343266 |
| gene-blatx_psbA_1 | 19032604,42 | -0,381971671 | 0,215182689 | -1,7751041 | 0,075880702 | 0,559846109 |
| gene-blatx_rpoC2_1 | 27566,16536 | 0,097390631 | 0,270134911 | 0,3605259 | 0,718453893 | 0,827783834 |
| gene-blatx_rpoC1_1 | 20580,2972 | 0,014303469 | 0,339146498 | 0,0421749 | 0,966359273 | 0,96885542 |
| gene-blatx_ndhI_1 | 24862,18968 | 0,172744366 | 0,196204439 | 0,88043047 | 0,378626157 | 0,658461475 |
| gene-blatx_rps7_1 | 51792,2717 | 0,4574474 | 0,216242448 | 2,11543758 | 0,03439267 | 0,394332022 |
| gene-blatx_psbC_1 | 816735,8857 | -0,143091172 | 0,143221682 | -0,9990888 | 0,317751697 | 0,653109223 |
| gene-blatx_rps11_1 | 18599,78172 | 0,160078405 | 0,187986703 | 0,8515411 | 0,39446884 | 0,663709477 |
| gene-blatx_rps8_1 | 9915,650207 | 0,315349225 | 0,267687096 | 1,17805165 | 0,238776017 | 0,637866774 |
| gene-blatx_atpE_1 | 81441,44932 | 0,275502364 | 0,186174369 | 1,47980823 | 0,138924431 | 0,613582901 |
| gene-blatx_ndhA_1 | 64267,13524 | 0,173428222 | 0,174286853 | 0,99507346 | 0,319700539 | 0,653109223 |
| gene-blatx_ndhC_1 | 28475,46761 | -0,167389434 | 0,31071289 | -0,538727 | 0,590075216 | 0,781849661 |
| gene-blatx_atpB_1 | 144294,5597 | 0,323162917 | 0,192467045 | 1,67905585 | 0,09314116 | 0,613582901 |
| gene-blatx_rbcL_1 | 5628759,686 | 0,155295168 | 0,130643751 | 1,1886919 | 0,23456093 | 0,637866774 |
| gene-blatx_ndhG_1 | 37225,405 | -0,126793653 | 0,145996816 | -0,8684686 | 0,385137844 | 0,658461475 |
| gene-blatx_rps15_1 | 6422,750703 | 0,27041804 | 0,191656999 | 1,41094789 | 0,158259979 | 0,637866774 |
| gene-blatx_rps16_1 | 85949,28409 | -0,281777158 | 0,209214538 | -1,3468335 | 0,178033848 | 0,637866774 |
| gene-blatx_rpoA_1 | 36067,94546 | 0,132568512 | 0,170369243 | 0,77812468 | 0,436495516 | 0,701038252 |
| gene-blatx_ndhD_1 | 72246,59474 | -0,095993405 | 0,246071841 | -0,3901032 | 0,69646026 | 0,820275417 |
| gene-blatx_psaC_1 | 147382,8502 | -0,253468438 | 0,279290839 | -0,907543 | 0,364119737 | 0,658461475 |
| gene-blatx_ccsA_1 | 56474,65699 | -0,310151517 | 0,251867155 | -1,2314091 | 0,218169879 | 0,637866774 |
| gene-blatx_cemA_1 | 48587,30946 | 0,201315129 | 0,216610176 | 0,92938907 | 0,352687493 | 0,658461475 |
| gene-blatx_ndhF_1 | 52342,28881 | -0,0750267 | 0,191610329 | -0,3915587 | 0,695384276 | 0,820275417 |
| gene-blatx_rps12_1 | 3963857,306 | -0,173761213 | 0,148105256 | -1,1732279 | 0,240704443 | 0,637866774 |
| gene-blatx_rpl14_1 | 12039,49383 | 0,352342609 | 0,28710699 | 1,2272171 | 0,219741003 | 0,637866774 |
| gene-blatx_rps3_1 | 20649,5716 | 0,389916716 | 0,28761669 | 1,35568181 | 0,175200428 | 0,637866774 |
| gene-blatx_accD_1 | 27277,08092 | -0,078235783 | 0,207557581 | -0,3769353 | 0,706221674 | 0,82263184 |
| gene-blatx_matK_1 | 331614,1921 | 0,450352869 | 0,216146915 | 2,08354983 | 0,037201134 | 0,394332022 |
| gene-blatx_petB_1 | 382149,002 | -0,114519989 | 0,19645776 | -0,5829242 | 0,559944305 | 0,775821811 |
| gene-blatx_rpl22_1 | 9994,041845 | 0,494690913 | 0,325517179 | 1,51970755 | 0,128584494 | 0,613582901 |
| gene-blatx_ycf1_1 | 41038,98652 | 0,049058444 | 0,147957719 | 0,3315707 | 0,740213451 | 0,843684148 |
| gene-blatx_rpl16_1 | 19159,49323 | 0,312866372 | 0,229865604 | 1,3610839 | 0,173487178 | 0,637866774 |
| gene-blatx_petL_1 | 21182,14921 | -0,760674041 | 0,244977556 | -3,1050765 | 0,001902298 | 0,100821819 |
| gene-blatx_infA_1 | 7172,550416 | 0,229830542 | 0,258725896 | 0,88831673 | 0,374370402 | 0,658461475 |
| gene-dPPRrpl2 | 118270,9705 | 15,53522742 | 0,490091457 | 31,6986293 | 1,6227E-220 | 1,7201E-218 |

**Table S3.** Differential gene expression analysis (DEseq2) data used for MA-plotting.

| Name | Sequence 5'-3' | Gene | Usage | Orientation |
|---|---|---|---|---|
| ***In planta* editing assay** | | | | |
| K1377 | TTACCGCAAGGCATAGAGGGGGAG | *rpl2* | RT-PCR - sequencing | For |
| K1378 | TGGCCGTGCCTAAGGGCATATCGG | *rpl2* | RT-PCR - sequencing | Rev |
| K1379 | TAGCTGCTTCAGCTTCAGCCAC | *ndhB* | RT-PCR - sequencing | For |
| K1380 | CAGTTCCGGTACGTAGACCAAATA | *ndhB* | RT-PCR - sequencing | Rev |
| K1413 | GGACCTCAACATCCTGCTGCT | *nad7* | RT-PCR - sequencing | For |
| K1415 | ATAGCATGAGTAGTTAAAGCAAGT | *nad7* | RT-PCR - sequencing | Rev |
| k177 | ACCCTTCCTCTATATAAGGAAG | *35S promoter* | genotyping- sequencing | For |
| K1233 | GTAGGAGCCACTCCCTGACG | *dPPRe* | genotyping- sequencing | Rev |
| K1396 | GCGAAAGAACGTCTGAAATTGGATCCG | *dPPRe* | genotyping- sequencing | For |
| K1382 | ACTTTGTACAAGAAAGCTGGGT | *attB2* | genotyping- sequencing | Rev |
| ***E. coli* editing assay** | | | | |
| PPR56-EDYWfor | GAAGCAACTTGGGGAG | *dPPRe* | RT-PCR | For |
| synPPRn113attB-for-long | TTGGTACCAAGCAGGCTtcccatcagacaccaac | *dPPRe* | cloning | For |
| rpl2_200bp_for | GTGGTGACTCGAATTTAAATgtctataccgtaaaatag | *rpl2* | cloning | For |
| rpl2_200bp_rev | gttGGCGCGccatttttataggaac | *rpl2* | cloning | Rev |
| ndhB_200bp_for | GTGGTGACTCGAATTTAAAtttcgatattcctttttatttc | *ndhB* | cloning | For |
| ndhB_200bp_rev | gttGGCGCGCCgctcgcatatccacc | *ndhB* | cloning | Rev |
| DYW_PPR_56_for | GAAATCCTATTCGTGTATTCAAGAACC | *dPPRe* | cloning | For |
| 169414DYW3rev | CGAAGGTTCTTGAATACACGAATAGG | *dPPRe* | cloning | Rev |
| petG_Smi_rev | AAATTCGAGTCACCAC | petG41Kmod | cloning | Rev |
| petG_30K40Kseqrev | GGTTATGCTAGTTATTGCTCAGC | petG41Kmod | RT-PCR- sequencing | Rev |

**Table S5.** Oligonucleotide sequences used in this study.
